# Amelioration of muscular dystrophy phenotype in mdx mice by inhibition of Flt1

**DOI:** 10.1101/609735

**Authors:** Mayank Verma, Yuko Shimizu-Motohashi, Yoko Asakura, James Ennen, Jennifer Bosco, Zhiwei Zou, Guo-Hua Fong, Serene Josiah, Dennis Keefe, Atsushi Asakura

## Abstract

Duchenne muscular dystrophy (DMD) is an X-linked recessive genetic disease in which the dystrophin coding for a membrane stabilizing protein is mutated. Recently, the vasculature has also shown to be perturbed in DMD and DMD model *mdx* mice. Data-mining DMD transcriptomics revealed the defects were correlated to a vascular endothelial growth factor (VEGF) signaling pathway. To reveal the relationship between DMD and VEGF signaling, *mdx* mice were crossed with constitutive (*CAG^/CreERTM^*:*Flt1^LoxP/LoxP^*) and endothelial cell-specific conditional gene knockout mice (*Cdh5^CreERT2^:Flt1^LoxP/LoxP^*) for *Flt1* which is a decoy receptor for VEGF. Previous work demonstrated that heterozygous global *Flt1* knockout mice increased vascular density and improved DMD phenotypes when crossed with DMD model *mdx* and *mdx:utrn^-/-^* mice. Here, we showed that while constitutive deletion of *Flt1* is detrimental to the skeletal muscle function, endothelial cell-specific *Flt1* deletion resulted in increased vascular density and improvement in the DMD-associated phenotype in the *mdx* mice. These decreases in pathology, including improved muscle histology and function, were recapitulated in *mdx* mice given anti-FLT1 peptides or monoclonal antibodies, which blocked VEGF-FLT1 binding. The histological and functional improvement of dystrophic muscle by FLT1 blockade provides a novel pharmacological strategy for the potential treatment of DMD.

## Introduction

Duchenne muscular dystrophy (DMD) is an X-linked muscle disease affecting one in 5,000 newborn males, in which the gene encoding the dystrophin protein is mutated. It is a progressive neurodegenerative disease with clinical symptoms manifesting at 2-3 years of age, loss of ambulation in early teen years and death by either respiratory insufficiency or cardiac failure in their 20s. A disease model for DMD is the *mdx* mouse, which lacks functional dystrophin expression due to a point mutation in the dystrophin gene. The *mdx* mouse has been extensively characterized and contributed to the understanding of the disease pathology (1).

Although the role of dystrophin in the skeletal muscle is widely appreciated, endothelium and vascular smooth muscle cells also express dystrophin (2). The absence of dystrophin in these cells induced vessel dilation and abnormal blood flow, resulting in a state of functional ischemia, worsening the muscle pathology in *mdx* mice (3). Restoration of dystrophin specifically in the smooth muscle of the vasculature rescued some aspects of the skeletal muscle pathology associated with the *mdx* mice (4). Disruption of the dystrophin-associated sarcoglycan complex in vascular smooth muscle perturbed vascular function resulting in exacerbation of muscular dystrophic changes (5). Dystrophin is responsible for anchoring neuronal nitric oxide synthase (nNOS) to the cell surface, which is crucial for exercise-induced increases in blood supply in muscle via NO-mediated vasodilation (6). Administration of a phosphodiesterase-5 (PDE-5) inhibitors to *mdx* mice, which increased NO production, rescued the muscle from this state of functional ischemia, and improved muscle function in *mdx* mice (7, 8). Similarly in humans, PDE-5 inhibitors given to both DMD boys and adult patients with Becker muscular dystrophy (BMD), a milder form of muscular dystrophy, alleviated functional ischemia during muscle contraction (9, 10). More recent data showed that *mdx* skeletal muscle was less perfused and displayed marked microvessel alterations compared to wild-type *C57BL6* mice (11, 12). While current studies support the importance of NO-mediated vasodilation in DMD, the relationship between DMD and angiogenesis is not well understood.

Vascular endothelial growth factor (VEGF) signaling is one of the strongest modulators of angiogenesis and includes the ligands VEGFA, VEGFB, VEGFC and PlGF. VEGFA is the most well studied ligand of the system and acts through its two receptors, VEGF receptor-1 (VEGFR1/FLT1) and VEGF receptor-2 (VEGFR2/FLK1/KDR). Although FLK1 possesses stronger signaling capabilities, FLT1 has considerably higher affinity for VEGF but weaker signaling capabilities. In normal tissue, FLT1 acts as a sink trap for VEGF thereby preventing excessive pathological angiogenesis. In addition, soluble FLT1 (sFLT1) functions as an endogenous VEGF trap (13). Despite the known angiogenic defect in DMD and *mdx* mice, it is not known whether VEGF and its receptors are implicated in this disease process. Previous data from our laboratory demonstrated that heterozygous *Flt1* gene knockout (*Flt1^+/-^*) mice were viable and displayed developmentally increased capillary density in the skeletal muscles (14). Importantly, when crossed *Flt1^+/-^* with *mdx* or *mdx:utrn^-/-^* mice, these mice displayed both histological and functional improvements of the dystrophic pathologic phenotype. However, it remained unknown whether postnatal *Flt1* gene deletion and pharmacological blockage of FLT1 could recapitulate these improvements in *mdx* mice.

In this report, we compared adult *mdx* mice with a constitutive conditional knockout and an endothelial cell-specific conditional knockout of *Flt1*. We showed that endothelial cell-specific *Flt1* deletion increased the capillary density in skeletal muscle and improved the DMD-associated muscle pathology. In addition, we showed that intravenous administration of anti-FLT1 peptides and monoclonal antibodies (MAbs) in *mdx* mice recapitulated the reduction in DMD-associated pathology seen after *Flt1* deletion in *mdx* mice, validating *Flt1* as a therapeutic target for the treatment of DMD.

## Results

### Postnatal *Flt1* gene deletion in mice display increase in capillary density

We previously found that *mdx* mice developmentally lacking one copy of the *Flt1* allele have increased muscle angiogenesis and improved muscle pathology (14). To investigate whether postnatal deletion of *Flt1* gene could affect the vasculature density, we crossed *CAG^CreERTM^* mice carrying a constitutively expressed *CreER^TM^* gene (15) with *Flt1^LoxP/LoxP^* mice (16) to generate conditional *Flt1* gene knockout (*CAG^CreERTM^:Flt1^LoxP/LoxP^*) mice (Figure 1A). Upon treatment with tamoxifen (TMX), which leads to global *Flt1* gene and FLT1 protein deletion (Supplemental Figure 1A, B), *Flt1^Δ/Δ^* mice displayed significantly increased CD31+ vascular density compared to the *Flt1^+/+^* (*Flt1^LoxP/LoxP^*) mice (Figure 1B, C). The increase in capillary density following TMX-mediated *Flt1* gene deletion was rapid (within 8 days) and long lasting (more than 207 days) (Figure 1D). This allowed us to be confident that we were able to phenotype late term changes following deletion of *Flt1* gene. Postnatal global loss of *Flt1* resulted in a reduction in body mass (17) without reduction of tibialis anterior (TA) muscle mass (Supplemental Figure 1C, D).

**Figure 1:**
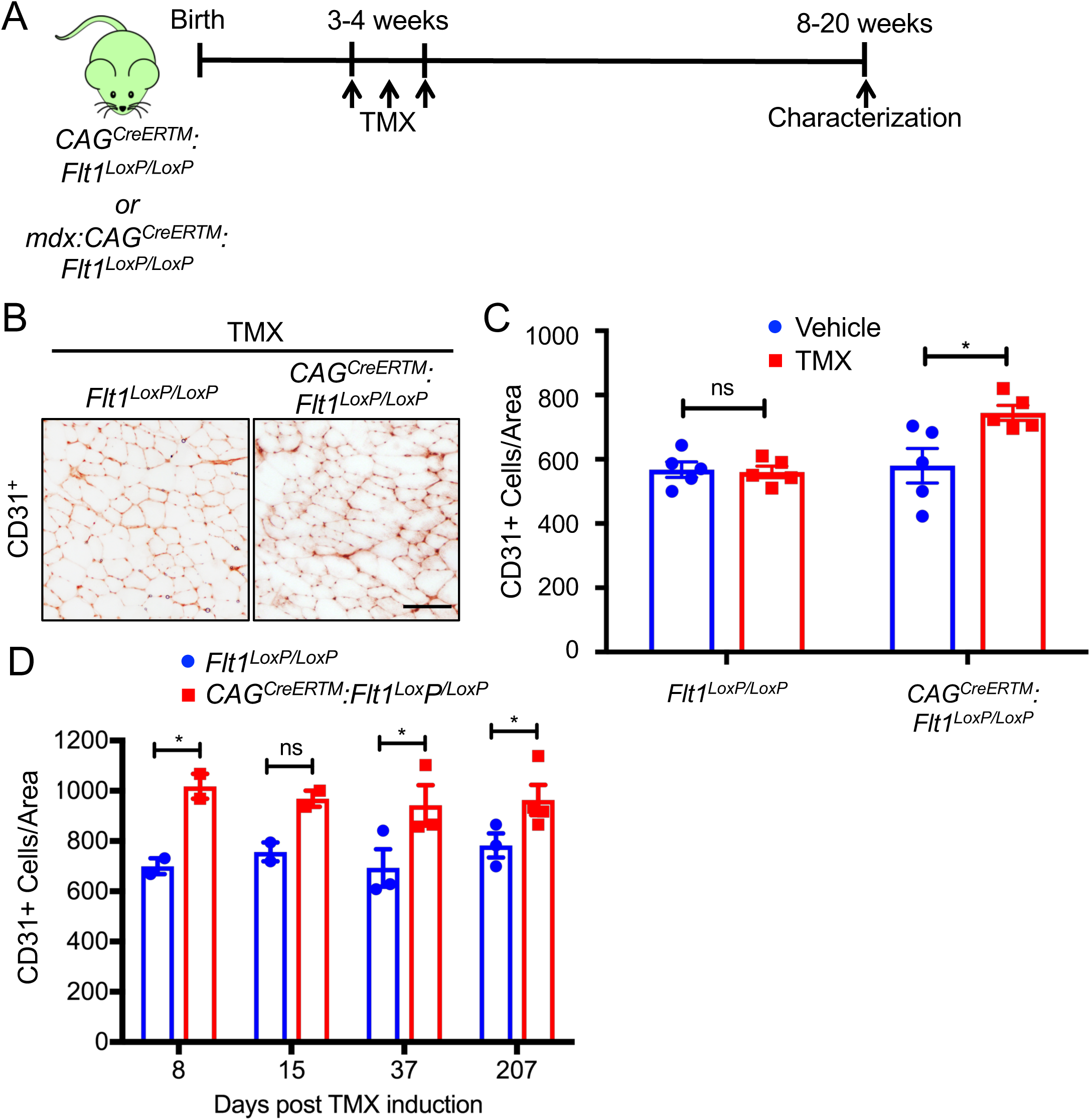
Postnatal deletion of *Flt1* can increase capillary density in skeletal muscle. A. Experimental scheme for assessing angiogenic response from conditional *Flt1* deletion. B. Representative images of CD31-stained cryosections from the skeletal muscle from *Flt1^Loxp/Loxp^* and *CAG^CreERTM^:Flt1^Loxp/Loxp^* mice following tamoxifen (TMX) induction. Scale bars indicate 100 µm. C. Increase in capillary density is dependent on *CAG^CreERTM^* as well as TMX induction in the *Flt1^Loxp/Loxp^* background. D. Increase in capillary density is rapid and sustained following TMX induction in *CAG^CreERTM^:Flt1^Loxp/Loxp^* mice more than 6 months following induction.

### *mdx:Flt1^Δ/Δ^* mice display worse muscle pathology compared to control *mdx:Flt1^+/+^* mice

As we did not see any gross changes in the skeletal muscle except for increased vascular density in the *Flt1^Δ/Δ^* mice, we crossed the *mdx* mice to the *CAG^CreERTM^:Flt1^LoxP/LoxP^* to obtain *mdx:CAG^CreERTM^:Flt1^LoxP/LoxP^* mice. We obtained these mice in expected mendelian ratios (Supplemental Figure 2A). Our original goal was to induce *Flt1* gene deletion prior to the onset of muscle pathology, thus before postnatal day 21 (p21), by treatment with TMX or its active form, 4-hydroxy tamoxifen (4-OHT). However, perinatal loss of *Flt1* resulted in lethality when TMX or 4-OHT treatment was initiated at p3 or p5 and partial lethality at p16 in *mdx:CAG^CreERTM^:Flt1^LoxP/LoxP^* (*mdx:Flt1^Δ/Δ^*) mice (Supplemental Figure 2B), indicating that *Flt1* is required in the perinatal stage for survival. The mice displayed no lethality when recombination was induced on or after p21. Importantly, the increase in capillary density by loss of the *Flt1* gene (Figure 1A) was maintained in the *mdx* background in the *mdx:Flt1^Δ/Δ^* mice (Figure 2A). This was accompanied by a physiological increase in skeletal muscle perfusion as shown by laser Doppler imaging (Figure 2B). The *mdx:Flt1^Δ/Δ^* mice showed a shift in fiber type composition toward increases in oxidative type I fibers (Supplemental Figure 3A, B). This was more pronounced in the EDL compared to the soleus, which is already predominantly type I.

**Figure 2:**
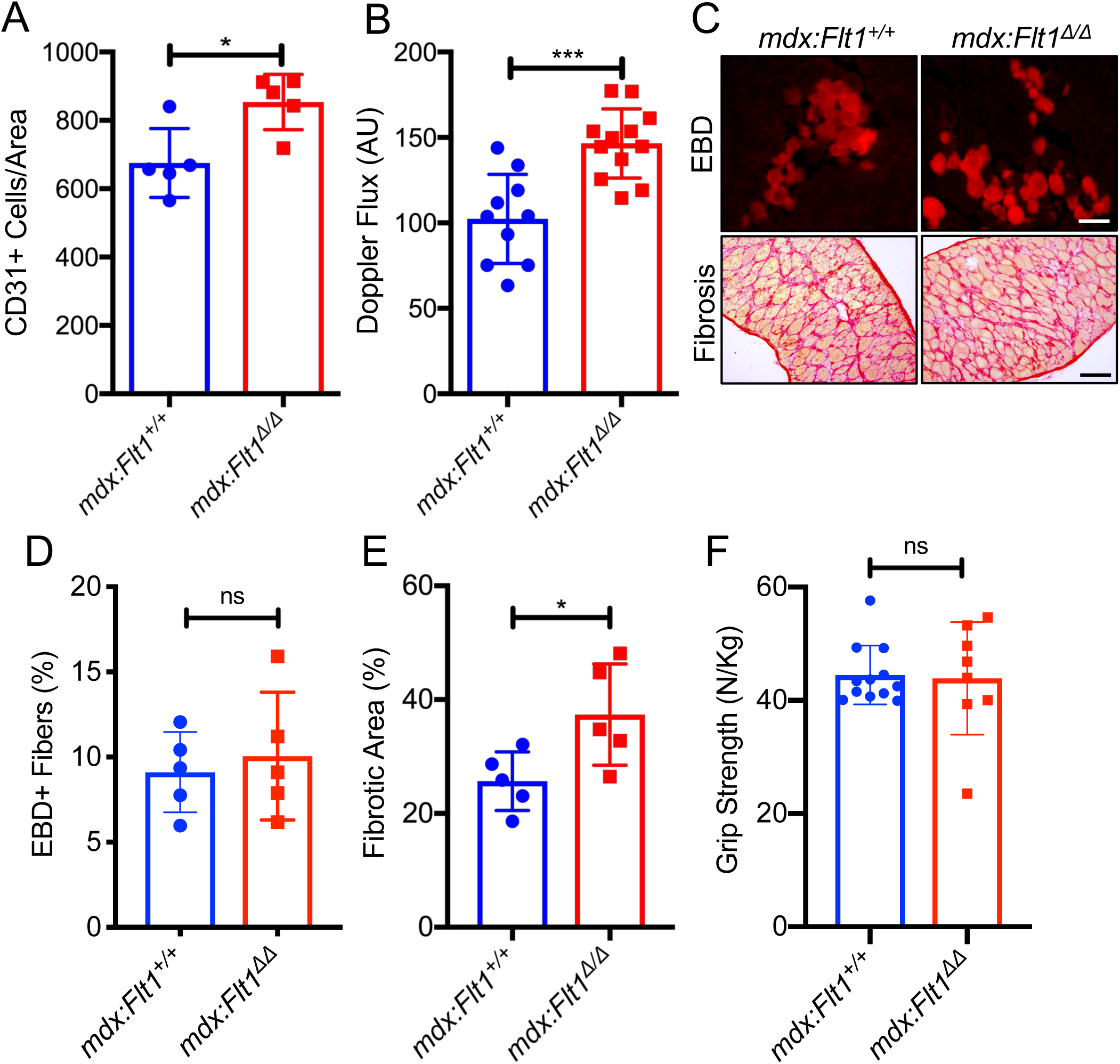
Increased angiogenesis is accompanied by worsened muscle pathology in the *mdx:Flt1^Δ/Δ^* mice. A. Capillary density is increased in *mdx:Flt1^Δ/Δ^* mice in the skeletal muscle. B. Skeletal muscle perfusion is increased in the TA muscle in *mdx:Flt1^Δ/Δ^* mice. C. Representative images of Evans blue dye (EBD) to measure acute damage and Sirius red staining to measure fibrosis in diaphragm. Scale bars indicate 100 µm. D. *mdx:Flt1^Δ/Δ^* mice show no difference in acute damage as judged by EBD. E. *mdx:Flt1^Δ/Δ^* mice show increased in fibrosis as evaluated by Sirius red staining. F. *mdx:Flt1^Δ/Δ^* mice show no difference in the grip strength normalized to body weight.

Next, we proceeded to look for dystrophinopathy-related muscle changes in the adult *mdx:Flt1^Δ/Δ^* mice. Similar to the *Flt1^Δ/Δ^* mice, body mass decreased without changing muscle mass in *mdx:Flt1^Δ/Δ^* mice compared with *mdx:Flt1^+/+^* mice (Supplemental Figure 3C, D). The *mdx:Flt1^Δ/Δ^* mice displayed notable white fur, a sign of premature aging or stress (Supplemental Figure 3E). Evans blue dye (EBD) accumulation (21) showed no differences, while fibrosis was markedly increased in the *mdx:Flt1^Δ/Δ^* mice (Figure 2 C, D, E). Embryonic MHC (eMHC)+ muscle fibers increased in the *mdx:Flt1^Δ/Δ^* mice (Supplemental Figure 3F), suggesting increased myofiber regeneration. There is no improvement in grip strength generation by the mice (Figure 2F). Taken together, the *mdx:Flt1^Δ/Δ^* mice showed an increase in capillary density but increased muscle pathology.

### Endothelial cell-specific loss of *Flt1* in *mdx* improves muscle phenotype in *mdx* mice

*Flt1* is expressed in several cell types including endothelial cells, myeloid cells and some neurons. Thus, we hypothesized that *Flt1* may be indispensable in one of these other compartments. Since endothelial cell-specific *Flt1* deletion resulted in increased capillary density in heart and adipose tissue (16, 22), we hypothesized that deletion of endothelial cell-specific *Flt1* would be sufficient to increase angiogenesis and improve muscle pathology in the *mdx* mice.

We attempted to increase the angiogenesis in skeletal muscle using an endothelial cell-specific *VE-cadherin (Cdh5)-CreERT2*-mediated *Flt1* deletion in mice. *Cdh5* is an endothelial cell-specific cadherin gene used for lineage tracing and conditional deletion of endothelial cells (23). Goel. et al. recently reported the presence of Cdh5 in satellite cells questioning the validity of using *Cdh5^CreERT2^* in skeletal muscle tissue (24). We verified the endothelial cell specificity of the *Rosa26R^mTmG^* reporter, and Cre-mediated excision resulted in the mGFP expression in the endothelial cells but no other cell types (Supplemental Figure 4A, B, C). We confirmed that the *Cdh5^CreERT2^* was not present in the Pax7+ satellite cells using single muscle fiber immunostaining (data not shown). We saw no difference in the body mass or muscle mass in endothelial cell-specific *Flt1* deleted mice compared with the control mice (Supplemental Figure 4D, E).

We crossed the *mdx:Flt1^LoxP/LoxP^* mice to the *Cdh5^CreERT2^* mice to yield the *mdx:Cdh5-Flt1^Δ/Δ^* mice (Figure 3A). Upon TMX treatment, capillary density and laser Doppler flow were increased in the skeletal muscle in *mdx:Cdh5-Flt1^Δ/Δ^* mice compared with *mdx:Cdh5-Flt1^+/+^* or *mdx:Cdh5-Flt1^+/Δ^* mice, indicating that endothelial cell-specific deletion of *Flt1* was sufficient to increase capillary density in the skeletal muscle (Figure 3B, C). This was accompanied by a physiological increase in skeletal muscle perfusion using laser Doppler (Figure 3D). Moreover, the *mdx:Cdh5-Flt1^Δ/Δ^* did not show any significant changes in body and muscle mass loss (Supplemental Figure 5). Signs of DMD-associated pathology such as increased EBD uptake and fibrosis were significantly reduced in the *mdx:Cdh5-Flt1^Δ/Δ^* mice (Figures 3B, E, F). The muscle fibers the *mdx:Cdh5-Flt1^Δ/Δ^* mice had decreased centrally located nuclei and maintained larger myofibers fibers compared with the *mdx:Cdh5-Flt1^+/+^* mice (Figure 4A, B, C). The *mdx:Cdh5-Flt1^Δ/Δ^* mice showed increased grip strength compared with the *mdx:Cdh5-Flt1^+/+^* mice (Figure 4D). Taken together, these data indicate that endothelial cell-specific *Flt1* loss was sufficient to increase capillary density and result in the histological improvements correlated with a functional improvement in the *mdx* mice.

**Figure 3:**
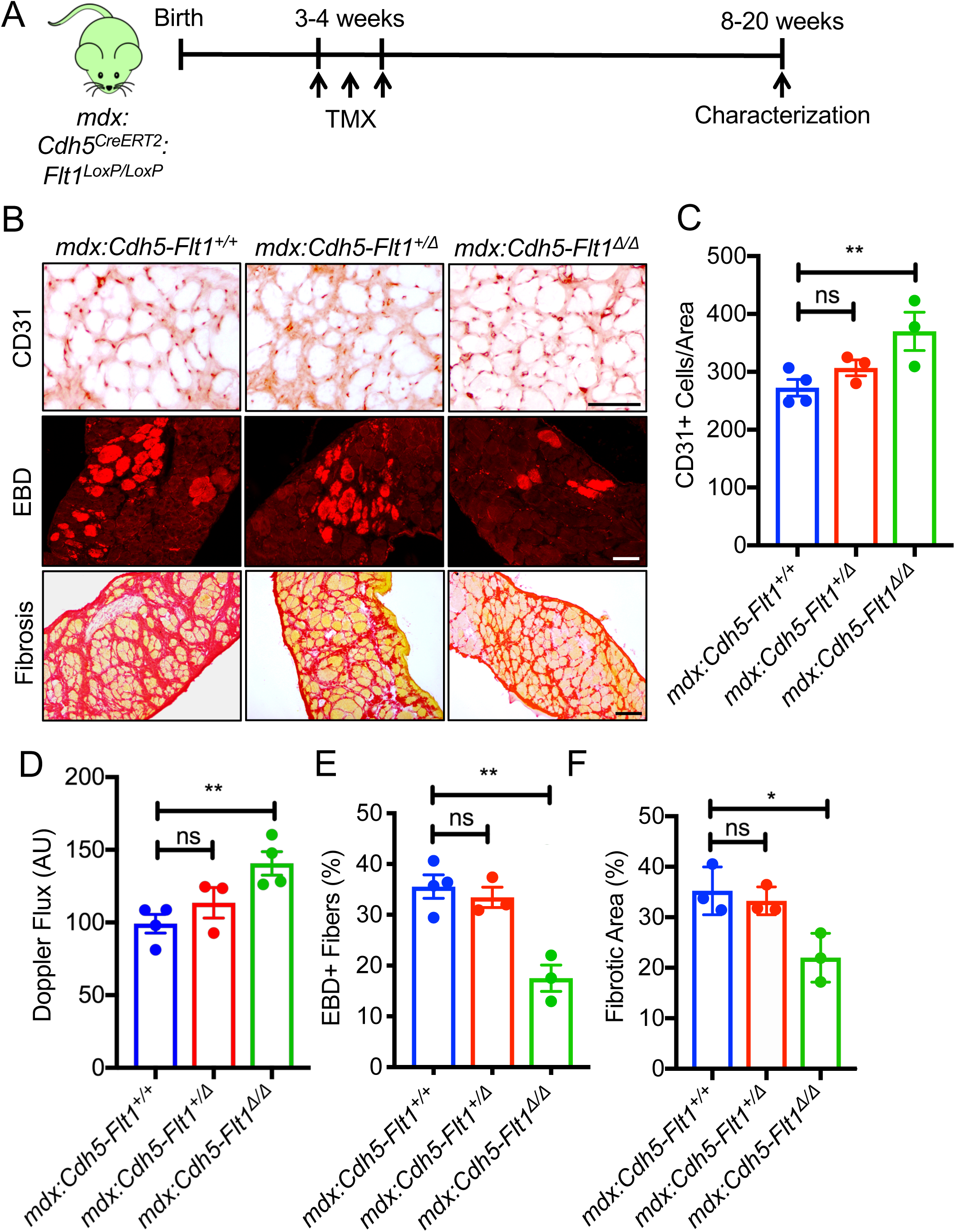
Endothelial cell-specific conditional deletion of *Flt1* in *mdx:Cdh5-Flt1^Δ/Δ^* mice improve capillary density and muscle phenotype. A. Experimental scheme for assessing angiogenic response from conditional *Flt1* deletion. B. Representative images of CD31 staining for capillary density, Sirius red staining to measure fibrosis and EBD to measure acute damage in the diaphragm. Scale bars indicate 100 µm. C. Endothelial cell specific conditional deletion of *Flt1* is sufficient to increase the capillary density in diaphragm. D. Endothelial cell specific conditional deletion of *Flt1* is sufficient to increase skeletal muscle perfusion. E. Acute damage as judged by EBD is reduced in *mdx:Cdh5-Flt1^Δ/Δ^* mouse muscle. F. Fibrosis is reduced in *mdx:Cdh5-Flt1^Δ/Δ^* mouse muscle as evaluated by Sirius red staining.

**Figure 4:**
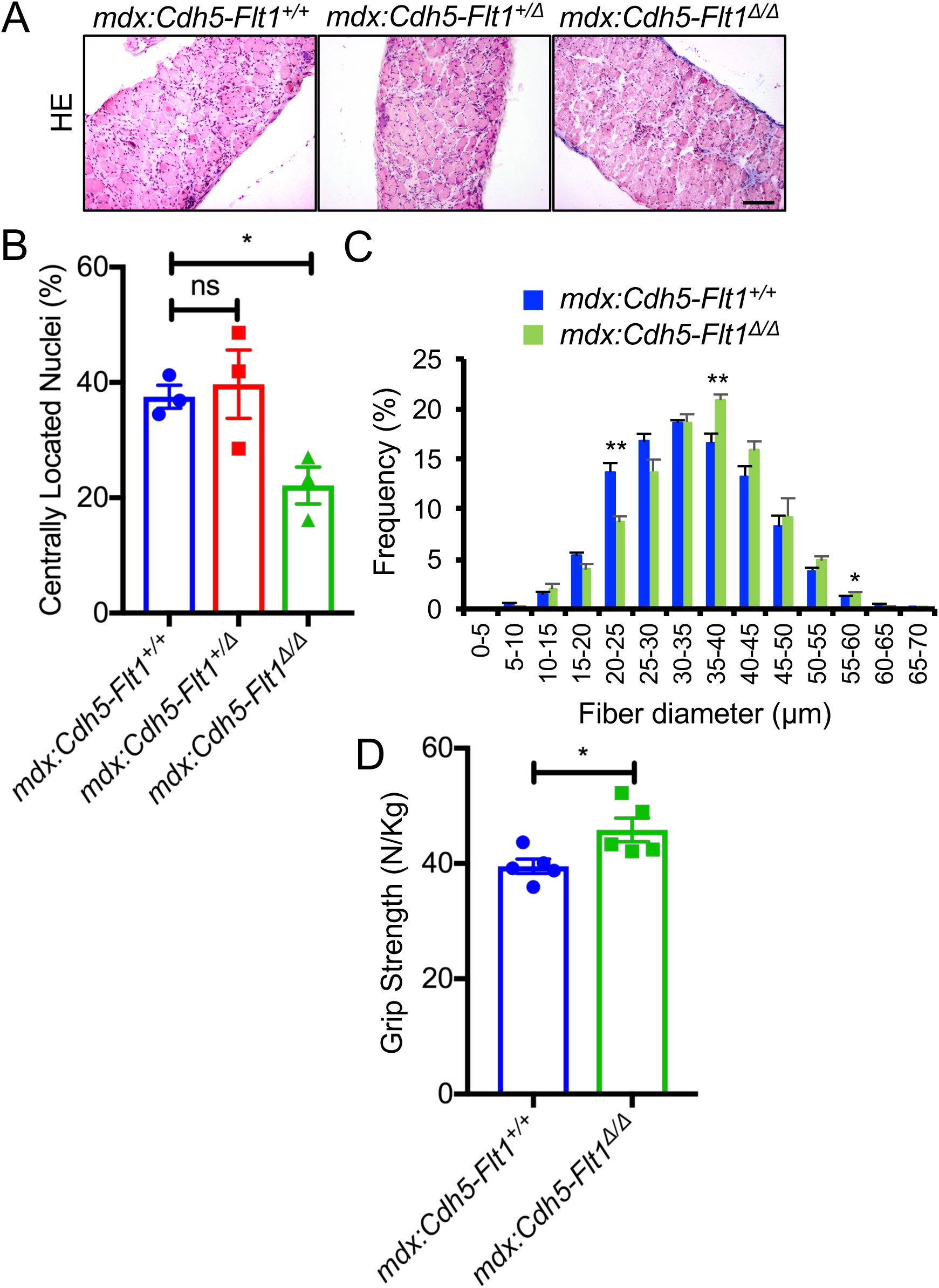
Endothelial cell-specific conditional deletion of *Flt1* in *mdx:Cdh5-Flt1^Δ/Δ^* mice improve muscle pathology. A. Representative images of HE staining of diaphragm. Scale bars indicate 100 µm. B. Diaphragm muscle fiber turnover is reduced in *mdx:Cdh5-Flt1^Δ/Δ^* mouse muscle as evaluated by centrally located nuclei (CLN). C. Distributions of mean fiber diameter in TA muscle of *mdx:Cdh5-Flt1^Δ/Δ^* mice were skewed toward the bigger fiber size compared with the control *mdx:Cdh5-Flt1^Δ/Δ^* mice. D. Grip strength is increased in *mdx:Cdh5-Flt1^Δ/Δ^* mice normalized to body weight.

### Pharmacological inhibition of FLT1 improved *mdx* mice

The genetic model of *Flt1* deletion showed an ameliorated phenotype in the *mdx* mice. To translate our genetic results into therapeutic approaches for DMD model mice, we utilized a previously reported anti-FLT1 hexapeptide (Gly-Asn-Gln-Trp-Phe-Ile or GNQWFI) that inhibits VEGF-binding to FLT1. Intramuscular administration of the anti-FLT1 peptide in TA muscle of perinatal *mdx* mice (Supplemental Figure 6A) increased capillary density and decreased muscle pathology in the treated muscle (Supplemental Figure 6B-D). We assessed the diaphragm muscle after systemic (IP) injection of the anti-FLT1 peptide at a low (10 mg/kg body weight) and a high dose (100 mg/kg body weight) to test for therapeutic potential in *mdx* mice (Figure 5A). While treatment with a low dose of anti-FLT1 peptide had no effect on capillary density, skeletal muscle perfusion, membrane permeability or fibrosis, the high dose increased capillary density, increased skeletal muscle perfusion, decreased EBD+ fibers, and decreased fibrosis (Figure 5B-F). Consequently, anti-FLT1 peptide-treated *mdx* mice increased grip strength compared with the control *mdx* mice (Figure 5F, G). There was no significant body mass alteration following anti-FLT1 peptide treatment (Supplemental Figure 6E, F). To increase stability and the hydrophobicity (25), we tested a D-isoform anti-FLT1 peptide attached to polyethylene glycol (PEG). However, systemic administration of the PEG-D-form anti-FLT1 peptide did not increase capillary density or improve muscle pathology or function in the *mdx* mice (Supplemental Figure 7). Taken together, these proof-of-concept experiments showed that postnatal inhibition of FLT1 by anti-FLT1 peptide could ameliorate the pathology associated with DMD in the *mdx* mice. However, the functional dose (100 mg/kg body weight) was orders of magnitude higher than the generally acceptable pharmacological standards of body weight dosage for small molecule drugs (26). A more potent or alternative strategy is required for further translational studies.

**Figure 5:**
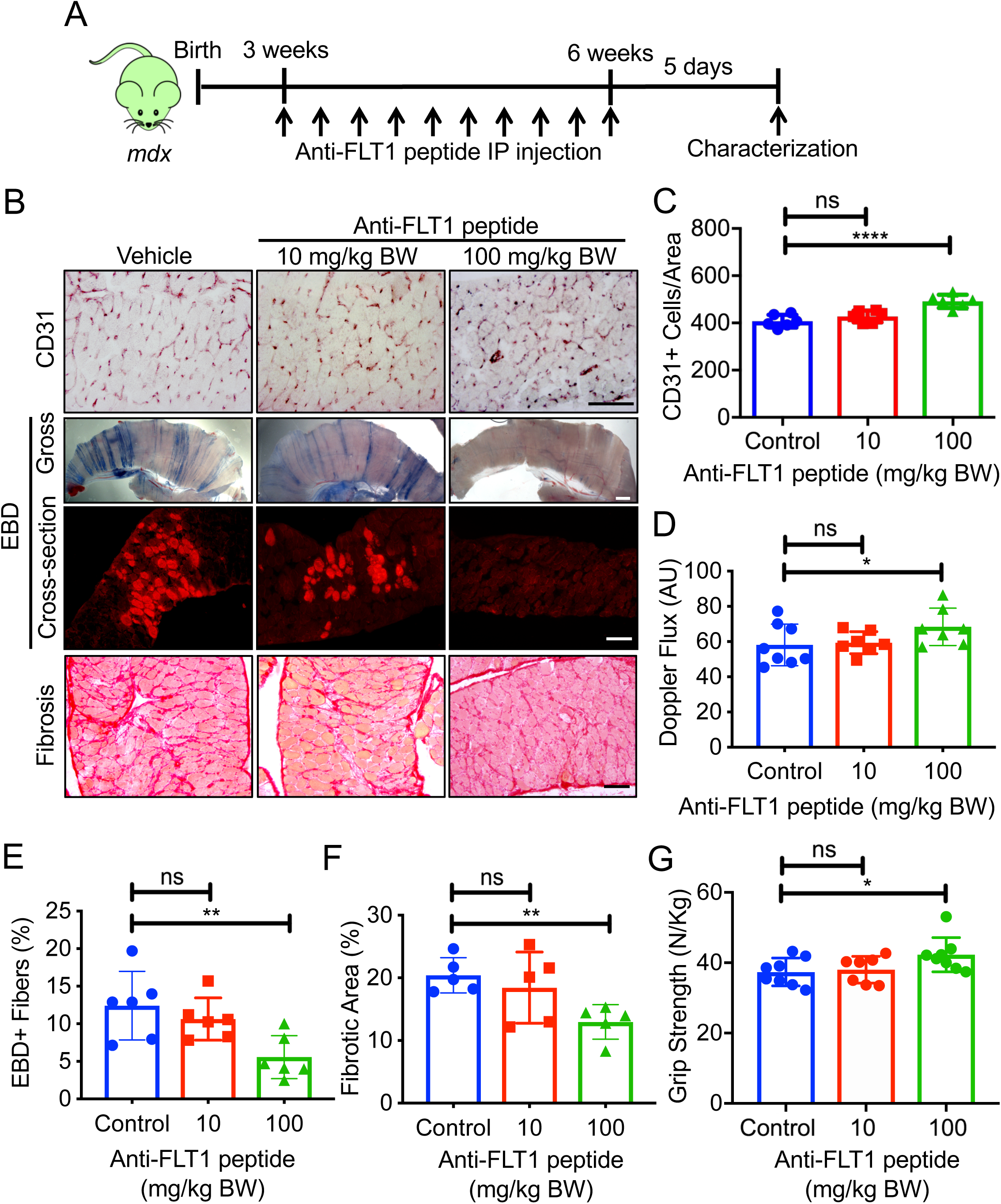
Systemic anti-FLT1 peptide improves skeletal muscle pathology in *mdx* mice. A. Experimental scheme for treatment of *mdx* mice with systemic treatment using anti-FLT1 peptide. B. Representative images of (top) CD31, (middle) gross whole mount and cryosections for EBD, (bottom) Sirius red staining of diaphragm in *mdx* mice treated with anti-FLT1 peptide. Scale bars indicate 100 µm. C. Anti-FLT1 peptide injection increases capillary density in the *mdx* mouse muscle at high dose. D. Anti-FLT1 peptide injection is sufficient to increase skeletal muscle perfusion in *mdx* mice at high dose. E. Anti-FLT1 peptide injection decreases EBD+ area in the *mdx* mouse muscle at high dose. F. Anti-FLT1 peptide injection decreases fibrotic area in the *mdx* mouse muscle at high dose. G. Grip strength is improved by anti-FLT1 peptide injection at high dose in *mdx* mice normalized to body weight.

### Screening for antibody against FLT1

As postnatal gene deletion and pharmacological inhibition of FLT1 decreased the muscular dystrophy-associated pathology in the *mdx* mice, we next sought to examine whether it could do this in a more translational manner using biologics to block FLT1. To establish proof of principle, we screened 8 commercially available MAbs that could block VEGF-FLT1 binding. We first screened for the ability of the MAbs to block chimeric FLT1-FC binding to PLGF2, a VEGF family protein, using ELISA (Supplemental Figure 8A, B), since both PLGF2 and VEGFA occupy the same binding sites on the extracellular domain of FLT1 (27). Three MAbs showed higher binding affinities: EWC (Novus Biologicals), MAB0702 Mab (Angio-Proteomie) and EWC (Acris GmbH) blocked binding by 65.4%, 64.8%, and 60.6%, respectively, while the control polyclonal anti-FLT1 antibody could block 83.1% of binding (Supplemental Figure 8B). We selected two MAbs (MAB0702 from Angio-Proteomie and EWC from Novus Biologicals) for further analyses based on their blocking efficiency and their availability for large *in vivo* studies. The MAB0702 and EWC were raised against the extracellular domains of the human FLT1, and likely target both membrane-bound (mFLT1) and sFLT1 due to similar homology of epitopes. We validated the blocking affinity to FLT1-VEGFA binding, and found that MAB0702 and EWC blocked binding by 40.1% and 19.9%, respectively, compared to the 94.3% for the polyclonal control (Supplemental Figure 8B). We further determined the antibodies’ affinities against mouse and human FLT1-FC protein using Biacore analysis (Supplemental Table 1). MAB0702 and EWC had similar association rate/binding constants for both mouse and human FLT1-FC. By contrast, MAB0702 had significantly higher dissociation constants for mouse FLT1-FC compared with EWC, while MAB0702 had lower dissociation constants for human FLT1-FC compared with EWC. Based on these affinity studies, we decided to utilize both MAB0702 and EWC MAbs for *in vivo* experiments.

### Testing Anti-FLT1 antibody treatment for DMD

We IV injected MAB0702 at a dose of 20 mg/kg body weight into *mdx* mice, and measured free sFLT1 and VEGFA in the serum. We found a significant decrease in free serum sFLT1 (Supplemental Figure 8C) and an increase in serum VEGFA levels (Supplemental Figure 8D) following MAB0702 injection, suggesting the efficient blocking of MAB0702 to sFLT1 and FLT1 *in vivo*. Based on our screening results, *mdx* mice were injected with IV MAbs or isotype control IgG dosing at 2 mg/kg or 20 mg/kg body weight every 3 days for four weeks beginning at 3 weeks of age (Figure 6A). While capillary density was not altered by treatment with either MAB0702 or EWC at 2 mg/kg body weight (data not shown), the treatment with MAB0702 at 20 mg/kg body weight but not EWC or isotype control significantly increased capillary density and skeletal muscle perfusion using laser Doppler (Figure 6B-D). More importantly, the mice treated with MAB0702 at 20 mg/kg body weight also displayed improved histology, such as decreased number of EBD+ fibers, decreased fibrosis, decreased calcification, and decreases in CLN (Figures 6B, E-G, 7A, B) without affecting any body weight compared with the *mdx* mice (Supplemental Figure 8E, F). Treatment prevented the increase in smaller caliber myofibers seen in the *mdx* mice (Figure 7A, C). While the EWC had significantly lower dissociation constants to mouse FLT1 compared with MAB0702 (Supplemental Table 1), we were surprised to find that *in vivo* administration of the EWC did not induce changes in capillary density, skeletal muscle perfusion or muscle pathology (Figures 6B, C-G, 7A, B). MAB0702-treated *mdx* mice generated increased grip strength compared with the *mdx* mice (Figure 7D). Skeletal muscle endurance, as assessed by treadmill running as an indicator of maximal muscle capacity, showed running duration and distance of MAB0702-treated *mdx* mice significantly increased compared to *mdx* mice (Figure 7E, F). Thus, anti-FLT1 antibody administration depleted free serum FLT1 levels and increased free serum VEGFA levels, which led to increased angiogenesis and reduced muscle pathology in *mdx* mice, providing a potential new pharmacological strategy for treatment of DMD.

**Figure 6:**
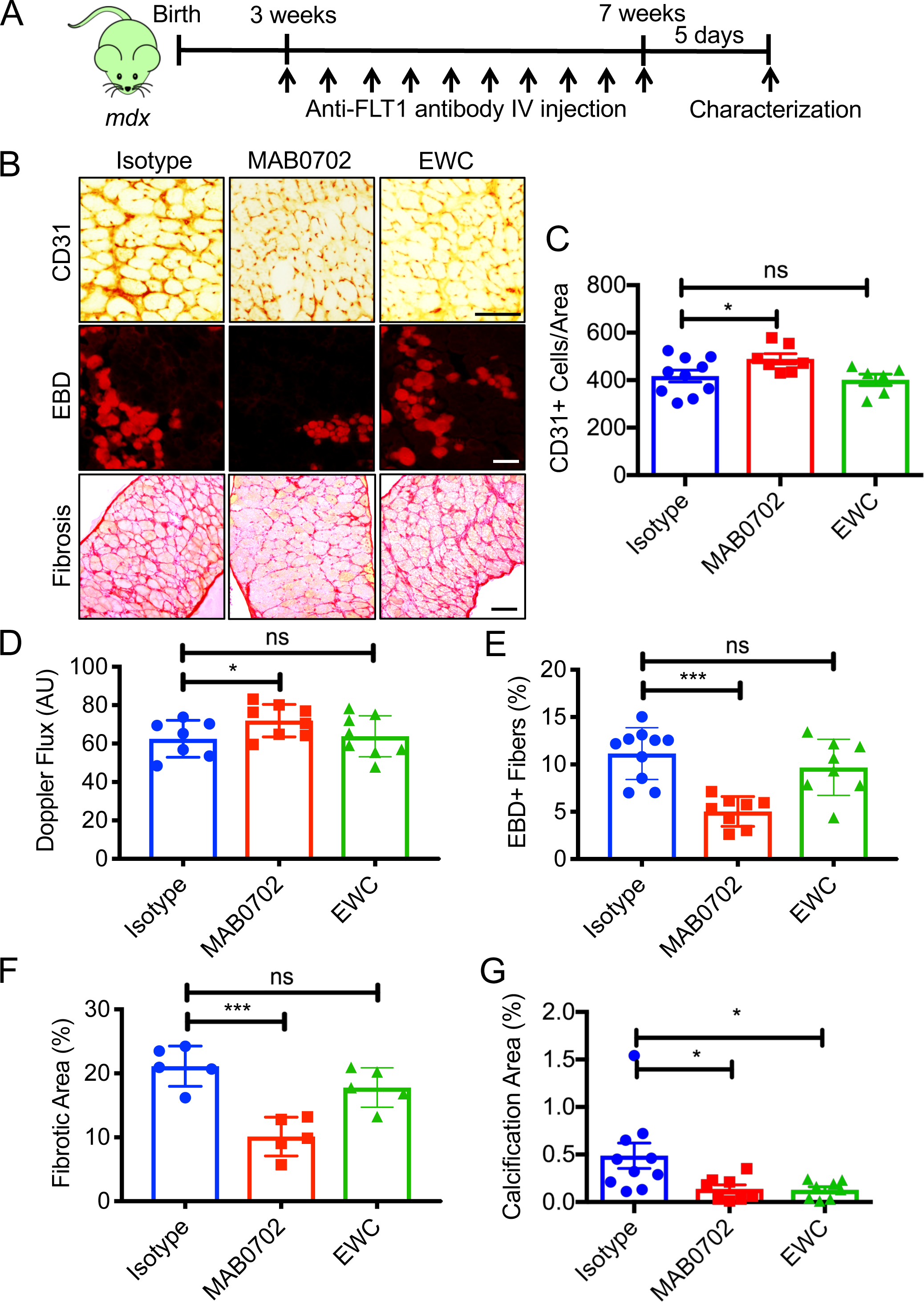
MAbs against FLT1 for the treatment of muscular dystrophy in the *mdx* mice. A. Experimental scheme for systemic treatment of *mdx* mice with systemic injection of anti-FLT1 antibody. B. Representative images of (top) CD31, (middle) EBD and (bottom) Sirius red staining of diaphragm in *mdx* mice treated with anti-FLT1 antibody. Scale bars indicate 100 µm. C. Capillary destiny is increased in MAB0702 antibody treated mice but not EWC antibody in *mdx* mice compared with isotype control. D. MAB0702 but not EWC injection is sufficient to increase skeletal muscle perfusion in *mdx* mice. E. EBD+ area is decreased in MAB0702 antibody treated mice but not EWC antibody in *mdx* mice compared with isotype control. F. Fibrotic area is decreased in MAB0702 antibody treated mice but not EWC antibody in *mdx* mice compared with isotype control. G. Calcification is decreased in MAB0702 antibody treated mice but not EWC antibody in *mdx* mice compared with isotype control.

**Figure 7:**
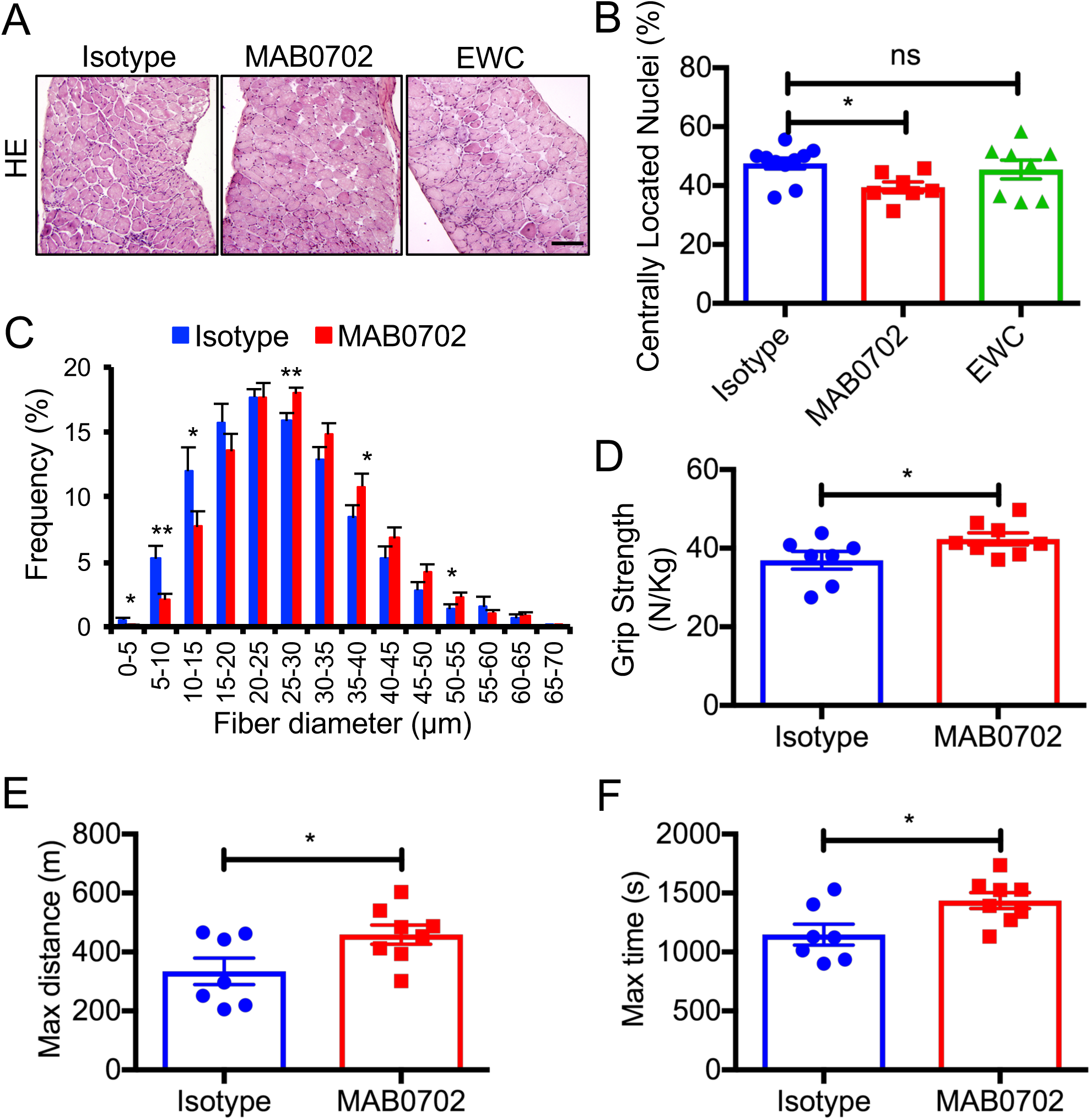
MAbs against FLT1 for the treatment of muscular dystrophy in the *mdx* mice. A. Representative images of HE staining of diaphragm in *mdx* mice treated with anti-FLT1 antibodies. Scale bars indicate 100 µm. B. Diaphragm muscle fiber turnover is reduced in MAB0702 treated *mdx* mouse muscle as evaluated by centrally located nuclei (CLN). C. Distributions of mean fiber diameter in TA muscle of MAB0702 treated *mdx* mice were skewed toward the bigger fiber size compared with compared with isotype control. D. Grip strength is improved in *mdx* mice treated with MAB0702 antibody compared with isotype control. E. Treadmill running time and distance (F) are improved in *mdx* mice following MAB0702 antibody treatment.

### VEGFA signaling is perturbed in DMD animal models and DMD patients

While angiogenic defects have been reported in the *mdx* mice, it is not known whether VEGF family and its downstream targets are implicated in dystrophinopathies. We probed the VEGF ligands and receptors in microarrays from skeletal muscles from *mdx* mice and the golden retriever muscular dystrophy (GRMD) canine model of DMD. *VEGFA* was downregulated in both models (Figure 8A). *Flt1* was downregulated in GRMD but not *mdx* muscles. To examine whether VEGF signaling is altered in DMD patients, we performed gene expression analysis on previously available data from microarrays and RNA-seq from patients with DMD. We also aggregated and probed microarray data from muscle biopsies of patients with various neuromuscular diseases or of healthy individuals after exercise. In the microarray data, *VEGFA* expression was increased after an acute bout of exercise, and VEGFA expression was reduced in ALS muscle, BMD muscle, as well as both early and late phases of DMD muscle (28) (Figure 8B). This was corroborated by RNA-seq data (Figure 8C). Angiogenic genes downstream of VEGF, such *HRAS*, *KRAS* and *NRAS,* were downregulated in DMD and BMD muscle despite an increase in VEGF pathways such as *HIF1A* (Figure 8D). By contrast, the expression of VEGF receptors (*Flt1* and *Flk1*) and the coreceptors (*Nrp1* and *Nrp2*) was not significantly changed in DMD muscle (Figure 8B, C). These data indicate that VEGFA expression is decreased in dystrophinopathy, and thus may benefit people with DMD by either increasing VEGFA and/or decreasing sFLT1 as a therapeutic target.

**Figure 8:**
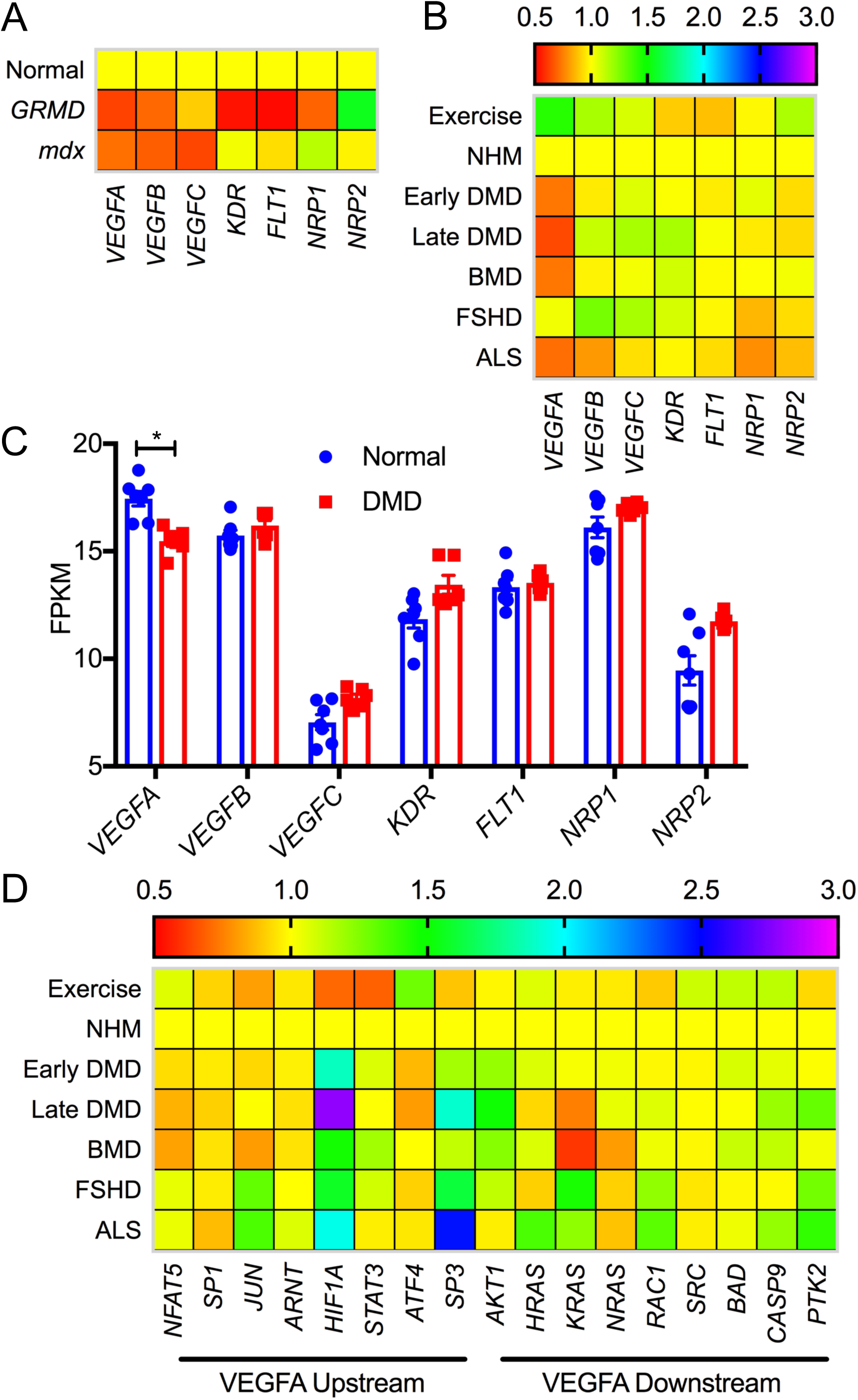
VEGFA expression is decreased in patients with DMD. A. *VEGFA* is compromised in both the dog (GRMD) and mouse (*mdx*) models of DMD compared with wild-type animals. B. Heatmap of VEGF ligand and receptor genes from microarrays from patients with neuromuscular diseases including early DMD (<age 3 yrs), late DMD, BMD, FSHD and ALS along with normal human muscle (NHM) and muscle following exercise. C. RNA-seq from patients with DMD show reduced *VEGFA* expression in muscle biopsies. D. Heatmap of genes upstream and downstream of *VEGFA* from microarray from patients with various neuromuscular diseases along with NHM and muscle following exercise.

## Discussion

In this report, we show that *Flt1* is important postnatally as conditional *Flt1* deletion in *mdx:Flt1^Δ/Δ^* mice in the perinatal stage results in lethality in mice. While deletion of *Flt1* in neonates increased capillary density in both C57Bl6 and *mdx* mice, it led to the worsening of the skeletal muscle phenotype. *Flt1* is expressed in several cell types including endothelial cells, myeloid cells and some neurons (23, 29–31). Thus, *Flt1* may be indispensable in one of these other compartments when perinatally deleted. For example, *Flt1* is expressed in motor neurons, where is not simply acting as a VEGFA sink-trap. In motor neurons, its tyrosine kinase activity it is responsible for their survival (29). Thus, the observed increased angiogenesis but worse pathological alterations in *mdx:Flt1^Δ/Δ^* muscle may be due to motor neuron–associated changes. By contrast, loss of *Flt1* in endothelial cells in the postnatal stage increased the capillary density and blood perfusion in skeletal muscle without significant decreases in weight or muscle mass. Increased angiogenesis and blood perfusion was observed in the global *Flt1* heterozygous knockout mice (14), indicating that loss of endothelial cell-specific *Flt1* in *mdx:Cdh5-Flt1^Δ/Δ^* mice was sufficient to produce increased capillary density and vascular perfusion in the skeletal muscle. This also led to an improved *mdx*-associate muscle pathology, confirming that postnatal deletion of *Flt1* is able to rescue the dystrophinopathy related muscle pathology.

Importantly, for the first time, we demonstrated that administration of both anti-FLT1 peptide and anti-FLT1 MAb increased angiogenesis, which led to an improved the pathology associated with DMD in *mdx* mice, and is a phenocopy of our genetic models (*mdx:Cdh5-Flt1^Δ/Δ^* mice). We screened commercially available MAbs for blocking ligands-FLT1, and demonstrated that administration of MAB0702 MAbs was able to phenocopy our genetic model in a manner suited for translational studies to reduce muscle pathology in *mdx* mice. Improvement of dystrophic muscle function by FLT1 blockade may provide a novel pharmacological strategy for treatment of diseases associated with DMD via increased serum and tissue VEGFA levels, which induce increased vascular density and blood perfusion.

After MAB0702 administration, we showed a small increase in VEGFA levels in serum. It should be noted that a mere 2-fold increase of VEGFA during development is incompatible with life in transgenic mice (32). Computation and experimental models showed that local VEGF gradients are more important than the total concentration (33–36). We recently demonstrated that muscle satellite cells express abundant VEGFA, which recruits endothelial cells and capillaries to the proximity of satellite cells on muscle fibers (37). Taken together, these data strongly suggest that hot-spots with high levels of VEGFA in *Flt1* knockout or anti-FLT1 treated mice may result in increased capillary density and vascular perfusion, which is predicted to result in decreased associated DMD-type pathological changes in the skeletal muscles.

While VEGFA binds to both FLT1 and FLK1, VEGFB, PlGF1 and PlGF2 only bind to FLT1 (38). This creates a scenario where PlGF1/2 and VEGFB binding can sequester FLT1, increasing serum and tissue VEGFA availability for VEGFA-FLK1 binding-mediated angiogenic induction. While *PlGF* is dispensable for normal development and health and not expressed in the normal adult tissues outside the female reproductive organs (39), VEGFB is expressed in the muscle tissue and muscle fibers (40). VEGFB may play a role in diet induced obesity and free fatty uptake by the endothelial cells in skeletal muscle (41), but this remains controversial (22). However, VEGFB overexpression does not result in an angiogenic response in ischemic skeletal muscle (42), indicating that blocking VEGFB-FLT1 signaling is not likely to be responsible for the angiogenic changes seen in this study.

The present study shows that increased angiogenesis may be a novel avenue to improve some of pathology associated with loss of dystrophin, including function, and could be used in conjunction with other treatment strategies. For example, intramuscular injection of VEGF containing recombinant adeno-associated viral vectors resulted in both functional and histological improvements in both ischemic and *mdx* muscle (19, 49, 50). This improvement was seen in combination with increased in angiogenesis in the muscle. Increased capillary density in the *mdx* muscle work through paracrine stimulation, by protecting muscle fiber damage and promoting satellite cell proliferation, survival and self-renewal in the vascular niche (37, 43). Christov et al. and our group showed that satellite cells were preferentially located next to capillaries (37, 44). Since our approaches increased the number of capillaries found in the examined muscles, it may effectively increase the amount of vascular niche that houses the satellite cell compartment in the muscle. Recently, we showed that genes encoding for endothelial and satellite cells were highly correlated across muscle groups, and endothelial cells could mediate satellite cell self-renewal via Notch activation (37). Therefore, increase in the vascular niche may increase satellite cell self-renewal and the number of myogenic precursor cells, which we hypothesize may be responsible for the improved phenotype seen in the *mdx* mice with *Flt1* deletion or functional blockage.

MAb-based therapeutics for the treatment of DMD have been developed to increase muscle mass via targeting myostatin and to decrease fibrosis via targeting fibroadipogenic progenitors (45). Further optimization using with humanized antibodies are required for future translation to humans (46). Taken together, we have gathered evidence for the first time that FLT1-targeted MAbs may be an effective therapeutic approach for the treatment of DMD.

## Methods

### Mice

*Flt1^Loxp/Loxp^* were obtained from Dr. Gua-Hua Fong (16). *Cdh5^CreERT2^* mice were obtained from Dr. Yoshiaki Kubota (23). B6Ros.Cg-*Dmd^mdx-5Cv^*/J (*mdx*; JAX stock #002379)(47), B6.Cg-Tg(CAG-cre/Esr1*)5Amc/J (*CAG^CreERTM^*; JAX stock #004682) (15). D2.129(Cg)-*Gt(ROSA)26Sor^tm4(ACTB-tdTomato,-EGFP)Luo^*/J (*Rosa26R^mTmG^*; JAX stock #007576) (48) were obtained from Jackson Laboratory. Colonies for all the mice were established in the laboratory. Cre recombination was induced using tamoxifen (TMX) (T5648, MilliporeSigma) dosed as 75 mg/kg body weight x 3 time over one week at 3-4 weeks of age unless otherwise specified. We also injected 4-hydroxy tamoxifen (4-OHT) (H6278, MilliporeSigma) dosed as 25 mg/kg body weight. Control mice contained the wild-type (WT) *CreER* allele or were injected with the vehicle (corn oil or 10% ethanol). All animal studies were approved by the IACUC at University of Minnesota.

### Anti-FLT1 peptides and antibodies

The anti-FLT1 peptide was synthesized from Peptide 2.0 Inc. based on the sequence (Gly-Asn-Gln-Trp-Phe-Ile or GNQWFI) as previously described (49). DMSO was used to dissolve the peptide. Twenty µg of peptide diluted in 2% DMSO in PBS solution for intramuscular injection per day in the TA muscle. Ten mg/kg body weight and 100 mg/kg body weight were used for the systemic treatment. The second generation anti-FLT1 peptide (PEG-G_D_N_D_Q_D_W_D_F_D_I_D_) was synthesized with the following modifications: The polyethylene glycol moiety was attached to the peptide to improve solubility in polar solvents and the D isomeric form was used instead of the L to enhance stability of the peptide (25). PBS was used as a vehicle. The peptides were commercially synthesized (LifeTein, LLC). Commercially available anti-FLT1 antibodies were obtained from the manufacturer in carrier free and preservative free form (AF471 from R&D Systems, D2 from Santa Cruz Biotechnology, EWC from Acris GmbH and Novus Biologicals, LS-C6855 from LifeSpan BioSciences, MAB1664 from MilliporeSigma and MAB7072 from Angio-Proteomie, Abcam and Santa Cruz Biotechnology). Isotype IgG (Santa Cruz Biotechnology) was used for control experiment. Two or 20 mg/kg body weight was used for the systemic treatment. Retro-orbital IV injections were performed for systemic treatment for both the peptides and the antibody treatment.

### RNA and genomic DNA isolation and qPCR

Mouse TA muscle was homogenized in TRIzol™ reagent (15596026, ThermoFisher Scientific) for RNA isolation. RNA was isolated using the Direct-zol™ RNA Microprep Kit (R2062, Zymo Research) with on-column DNase digestion followed by cDNA synthesis using the Transcriptor First Strand cDNA synthesis kit (04379012001, Roche Molecular Diagnostics) using random primers. Genomic DNA was isolated from mouse tail snips with lysis buffer containing Protenase K (P2308, MilliporeSigma). Genotyping was performed by agarose gel electrophoresis-mediated detection following PCR reaction by Taq polymerase (M0273, New England Biolabs). qPCR was performed using GoTaq® *qPCR* Master Mix (A6001, Promega). Primer sequences are listed in Table S1. All primers were synthesized as custom DNA oligos from Integrated DNA technologies (IDT).

### Muscle perfusion

RBC flux was evaluated using the moorLabTM laser Doppler flow meter as previously described (14). with the MP7a probe that allows for collecting light from a deeper tissue level than standard probes according to the manufacturer’s instructions (Moor Instruments). The fur from the right hind leg was removed using a chemical depilatory. Readings were taken using the probe from at least 10 different spots on the TA muscle. The AU was determined as the average AU value during a plateau phase of each measurement.

### Grip strength test

Forelimb grip strength test was performed following a previously published procedure (50). Briefly, *mdx* mice were gently pulled by the tail after fore limb-grasping a metal bar attached to a force transducer (Columbus Instruments). Grip strength tests were performed by the same blinded examiner. Five consecutive grip strength tests were recorded, and then mice were returned to the cage for a resting period of 20 minutes. Then, three series of pulls were performed each followed by 20 min resting period. The average of the three highest values out of the 15 values collected was normalized to the body weight for comparison.

### Treadmill running

Exer-3/6 Treadmill (Columbus Instruments) was used for treadmill running test as previously described (51). Briefly, for acclimation, mice were placed in each lane and forced to run on a treadmill for 5 minutes at a speed of 10 m/min on a 0% uphill grade for 3 days. And then, mice were forced to run on a treadmill with a 10% uphill grade starting at a speed of 10 m/min for 5 minutes. Every subsequent 2 minutes, the speed was increased by 2 m/min until the mice underwent exhaustion which was defined as the inability of the mice to remain on the treadmill. The time of running as well as the distance run were recorded.

### Histology and immunofluorescence

To assess the microvasculature ECs, we utilized intravital tomato-lectin (DL-1178, Vector labs) staining following retro-orbital IV injections. Tissues were frozen fresh using LiN_2_ chilled isopentane and stored at −80°C. Eight µm thick transverse cryosections were used for all histological analysis. Slides were fixed using 2% PFA and washed twice using PBS + 0.01% Triton (PBST) before being used. For capillary density measurement, immunohistochemistry for CD31 was performed using anti-CD31 antibody followed by the Vectastain Elite ABC Kit (PK-6100, Vector Laboratories) according to the manufacturer’s instructions and developed using 3-amino-9-ethylcarbazole (AEC) (A5754, MilliporeSigma). For immunofluorescence, sections were fixed, washed twice in PBST and blocked 5% goat serum and 0.2% Triton X-100 for 30 min. Sections were incubated with primary antibodies overnight, washed twice in PBST and incubated in fluorescent-conjugated secondary antibody for 1 hour. Anti-FLT1 (RB-9049, NeoMarkers) and anti-Laminin (4H8-2, MilliporeSigma) antibodies followed by anti-rabbit Alex-488 and anti-rat Alexa-568 (Molecular Probes) were used for detection of FLT1 expression in capillaries. Anti-slow MHC (NOQ7.5.4D, MilliporeSigma) or anti-embryonic MHC (F1.652, Developmental Study Hybridoma Bank) and anti-Laminin (4H8-2, MilliporeSigma) antibodies followed by anti-mouse Alex-488 (A21202, ThermoFisher Scientific) and anti-rat Alexa-594 antibodies (A21471, ThermoFisher Scientific) were used for detection of slow muscle fibers. One percent Evans blue dye (EBD) (E2129, MilliporeSigma) dissolved in PBS were injected IP at 1% body weight 16-20 hours prior to dissecting the mouse (21). Sirius red, Alizarin red, and Hematoxylin & Eosin (HE) were performed as previously described (14). All sections were co-stained using DAPI and mounted using DAKO mounting media. Microscopic images were captured by a DP-1 digital camera attached to BX51 fluorescence microscope with 10x, 20× or 40× UPlanFLN objectives (all from Olympus). Fiji was used for image processing (52).

### Biacore Surface Plasmon Resonance (SPR) Binding Assay for Anti-FLT1 MAbs

The single cycle kinetics method was used for sFLT1 binding assay by Biacore (GE Healthcare Bio-Science). A CM5 series S sensor chip (GE Healthcare Bio-Science) with mouse and human soluble FLT1-FC chimeric protein (471-F1-100 and 321-FL-050/CF, R&D Systems) immobilized to about 1,000 RU was used as ligand for the analyte binding of anti-FLT1 MAbs. An analyte range of 0-5 nM was used for kinetic experiments. A 5 min association step was used for each dilution followed by a 40 min dissociation. Chip surface was regenerated using pH 2.0 glycine between experiments.

### ELISA

For the initial FLT1 blocking MAb screening, Nunc 96 well ELISA plates (44-2404-21, ThermoFisher Scientific) were incubated with mouse FLT1-FC chimeric protein with His-tag (471-F1-100, R&D Systems). Plates were washed with 0.05% Tween 20/PBS, and blocked with 5% BSA (BP1600-1, Fisher Scientific)/PBS. After washing, diluted anti-FLT1 antibodies were incubated. After washing, recombinant mouse PlGF2 (465-PL-010/CF, R&D Systems) was incubated. After washing, biotin-conjugated anti-PlGF2 antibody (BAF465, R&D Systems) was added followed by a Streptavidin-HRP (R&D Systems, DY998). For a colormetric detection, plates were developed with 3,3′,5,5′-Tetramethylbenzidine (TMB) Liquid Substrate (T0440, MilliporeSigma) and the reaction was stopped with 0.5M H_2_SO_4_. Specs were read at 450 nm on spectramax M5 plate reader (Molecular Devices). For VEGFA-FLT1 blocking assays, 96 wells ELISA plates were incubated with recombinant mouse VEGFA (VEGF_164_) (493-MV-025/CF, R&D Systems,). After washing, anti-FLT1 antibodies were incubated with mouse FLT1-FC chimeric protein with His-tag (471-F1-100, R&D Systems). After incubation the antibody-FLT1-FC complex was added to the VEGFA-coated plates. A biotin conjugated anti-His-tag antibody (NB100-63172, Bio-Connect) was added followed by a Streptavidin-HRP (DY998, R&D Systems). For a colormetric detection, TMB was used as described above. For measurement of free sFlt1 and VEGFA in the serum. Animals were IV injected with 20 mg/kg body weight either isotype control IgG or the MAB0702 twice a weekly for four weeks beginning at 4 weeks of age. Blood was collected for biomaker analysis. Serum concentrations of sFLT1 and VEGFA were measured by ELSA kits following company recommended protocols (DY471 and DY493, R&D Systems). For a colormetric detection, TMB was used as described above.

### Microarray and RNA-seq analysis

Microarray analysis was performed using the Affymetrix Transcriptome Analysis Console (TAC). Samples in each experiment were RMA normalized and the expression was acquired using the Affeymetrix Expression analysis console with gene level expression. Heatmaps were generated in the Graphpad 7.1 (Prism). The code for generating each graph is listed in the following table, along with the link to the data in tabular format. All the data was obtained from NCBI GEO: Exercise, ALS, DMD, BMD, FSHD GSE3307, Early DMD GSE465, *mdx* GSE466, GRMD GSE69040, Satellite cells GSE15155. All arrays were normalized to their respective controls.

### Statistics

Statistics and graphs were calculated using Graphpad 7.1(Prism). Students t-test or ANOVA was used to compare two or more groups. Multiple comparison adjustment was performed with comparisons of 3 groups or more. * indicates p<0.05, ** indicates p<0.01, *** indicates p<0.001, *** indicates p<0.0001.

## Author contributions

Designing research study- MV, AA. Conducting experiments- MV, YM, JE, acquiring data- MV, YM, analyzing data- MV, YM, providing reagents- DK, SJ, initial antibody screening process- JB, ZZ, and writing manuscript- MV, AA. All the authors approved the final manuscript.

## Acknowledgments

We would like to thank Jake Trask for a critical reading. This work was supported by NIHT32-GM008244 and NIHF30AR066454 to MV, and NIH R03, MDA and a grant from Shire Human Genetic Therapies Inc., a member of the Takeda group of companies to AA.

## Supplemental Information

**Supplemental Figure 1: Confirmation of conditional deletion of *Flt1***

A. Immunostaining for FLT1 (green, arrows) and Laminin (red) shows effective deletion of FLT1 in TA muscle of *Flt1^Δ/Δ^* mice following tamoxifen (TMX) injection. DAPI staining (blue) is for all nuclei. Scale bars indicate 20 µm.

B. RT-qPCR shows deletion of the *Flt1* exon 3 in the TA muscle of *Flt1^Δ/Δ^* mice while exons 1 and 2 are retained.

C. *Flt1^Δ/Δ^* male mice show no difference in the body mass during the time course or (D) TA muscle compared to *Flt1^+/+^* male mice.

**Supplemental Figure 2: Generation of *mdx:CAG^CreERTM^:Flt1^Loxp/Loxp^* mice**

A. *mdx:CAG^CreERTM^:Flt1^Loxp/Loxp^* breed with *mdx:Flt1^Loxp/Loxp^* mice, and yield mice in expected ratios in a total of 92 mice genotyped. Chi-squared test shows no difference in expected.

B. Induction of *CreER^TM^* by TMX or 4-hydroxy-tamoxifen (4-OHT) shows that *Flt1* deletion in *mdx* mice prior to p21 results in partial or complete lethality in the *mdx:Flt1^Δ/Δ^* but not control *Flt1^+/+^* mice.

**Supplemental Figure 3: Phenotyping of *mdx:Flt1^Δ/Δ^* mice**

A. Representative image of slow MHC (green) and Laminin (red) in EDL and soleus muscle in *mdx:Flt1^Δ/Δ^* mice. Scale bars indicate 200 µm.

B. Slow MHC+ fibers are increased in the *mdx:Flt1^Δ/Δ^* mice in both EDL and Soleus muscle.

C. *mdx:Flt1^Δ/Δ^* mice show reduced male body mass but no difference in (D) muscle mass.

E. *mdx:Flt1^Δ/Δ^* mice show signs of premature aging such as white hair.

F. eMHC staining shows decreased fiber stability in muscle fibers in the soleus but not the EDL muscle in the *mdx:Flt1^Δ/Δ^* mice.

**Supplemental Figure 4: Validation of *Cdh5^CreERT2^* and *mdx:Cdh5-Flt1^Δ/Δ^* mice**

A. Experimental scheme for assessing angiogenic response from conditional *Flt1* deletion.

B. *Cdh5^CreERT2^:Rosa26R^mTmG^* mice reveal that *Cdh5^CreERT2^* efficiently induced mGFP expression in the capillaries (green) labeled with lectin (purple), but not in other cell types including muscle fibers (red). Scale bars indicate 20 µm.

C. *Cdh5^CreERT2^:Rosa26R^mTmG^* mice show efficient mGFP labeling of lectin+ and CD31+ endothelial cells. Histogram indicates that more than 90% of the cells are CD31+mGFP+ or lectin+mGFP+.

D. Body mass and (E) TA muscle mass are unchanged in *Cdh5-Flt1^+/+^* and *Cdh5-Flt1^Δ/Δ^* mice.

**Supplemental Figure 5: Validation of *mdx:Cdh5-Flt1^Δ/Δ^* mice**

A. Body mass and (B) TA muscle mass are unchanged in *mdx:Cdh5-Flt1^+/+^, mdx:Cdh5-Flt1^+/Δ^ and mdx:Cdh5-Flt1^Δ/Δ^* mice.

C. D. Body mass is unchanged in male or female *mdx:Cdh5-Flt1^+/+^* and *mdx:Cdh5-Flt1^Δ/Δ^* mice during the time course.

**Supplemental Figure 6: Efficacy of anti-FLT1 peptide following intramuscular injection in the TA muscle of *mdx* mice**

A. Experimental scheme for proof of principle study of *mdx* mice with intramuscular injection of anti-FLT1 peptide.

B. Representative images of CD31 (top) and EBD staining (bottom) of TA muscle injected with anti-FLT1 peptide. Scale bars indicate 50 µm.

C. Neonatal intramuscular injection of anti-FLT1 peptide increases capillary density in the TA muscle of the *mdx* mice.

D. Neonatal intramuscular injection of anti-FLT1 peptide decreases EBD+ area in the TA muscle of the *mdx* mice.

E. F. Systemic anti-FLT1 peptide injection does not change body mass in the male or female *mdx* mice at low or high dose.

**Supplemental Figure 7: Systemic PEG-anti-FLT1 peptide injection does not improve skeletal muscle pathology in *mdx* mice**

A. Experimental scheme for systemic treatment of *mdx* mice with IP injection of PEG-anti-FLT1 peptide.

B. Systemic PEG-anti-FLT1 peptide injection does not change body mass in the male *mdx* mice at low or high dose.

C. Systemic PEG-anti-FLT1 peptide injection does not increase capillary density in the *mdx* mice at low or high dose.

D. Systemic PEG-anti-FLT1 peptide injection does not decrease EBD in the *mdx* mice at low or high dose.

E. Systemic PEG-anti-FLT1 peptide injection does not improve grip strength in the *mdx* mice at low or high dose.

**Supplemental Figure 8: Screening for commercially available antibodies against FLT1 and phenotyping of *mdx* mice treated with MAB0702**

A. Commercially available MAbs for anti-FLT1 screened for blocking activity against PlGF using ELISA. AF471 polyclonal anti-FLT1 antibody was used as a positive control. AP, Angio-Proteomie; SC, Santa Cruz Biotechnology.

B. Two selected MAbs screened for blocking activity against VEGFA using ELISA. AF471 polyclonal anti-FLT1 antibody was used as a positive control. AP, Angio-Proteomie.

C. Serum free sFLT1 is decreased in mice injected with MAB0702 compared to isotype control.

D. Serum free VEGFA is increased following MAB0702 treatment.

E. F. Systemic anti-FLT1 antibody injection does not change body mass in the male or female *mdx* mice.

