## Supplemental Figures for "Amelioration of muscular dystrophy phenotype in mdx mice by inhibition of Flt1"

Figure S1

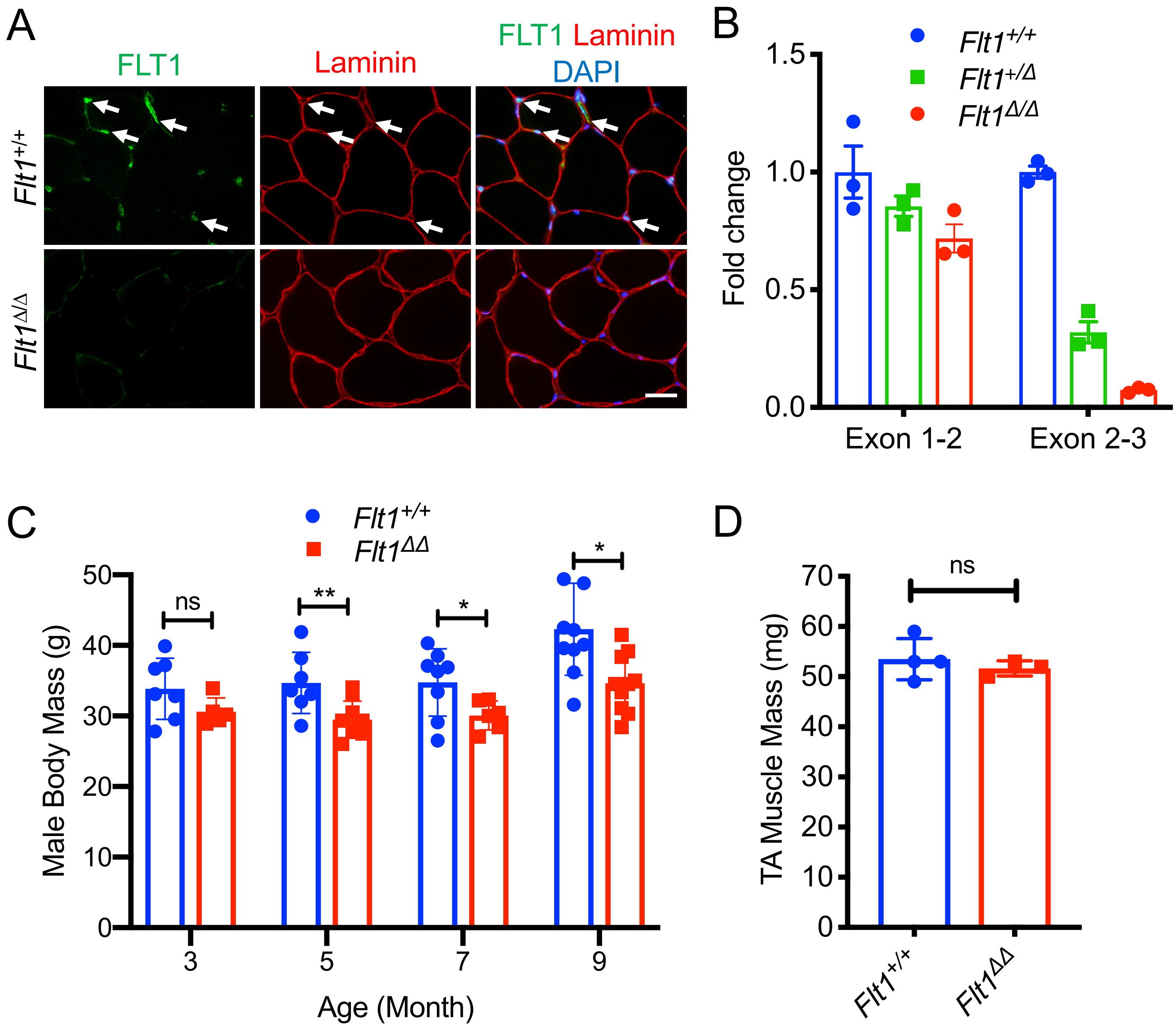

A
FLT1
Laminin
FLT1 Laminin DAPI
Flt1+/+
Flt1Δ/Δ
B
Flt1+/+
Flt1+/Δ
Flt1Δ/Δ
Fold change
1.5
1.0
0.5
0.0
Exon 1-2
Exon 2-3
C
Flt1+/+
Flt1ΔΔ
Male Body Mass (g)
50
40
30
20
10
0
ns
**
*
*
3
5
7
9
Age (Month)
D
TA Muscle Mass (mg)
70
60
50
40
30
20
10
0
ns
Flt1+/+
Flt1ΔΔ

Figure S2

A

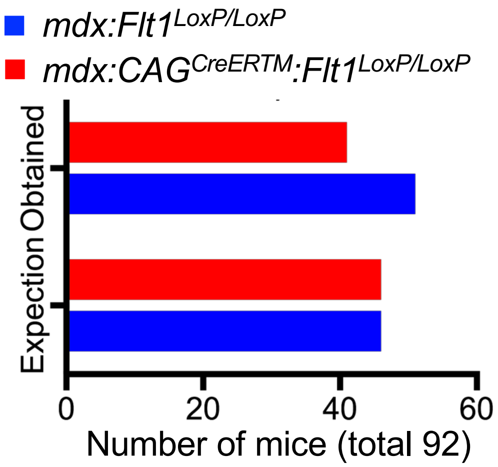

B

| | Viability 5 days Post Tamoxifen Injection | | | |
|---|---|---|---|---|
| | *mdx:Flt1*$^{LoxP/LoxP}$ | | *mdx:CAG*$^{CreERTM}$*:Flt1*$^{LoxP/LoxP}$ | |
| Tamoxifen Start Age | Vehicle | TMX or 4-OHT | Vehicle | TMX or 4-OHT |
| Postnatal day 3 (p3) | N/A | 12/14 (TMX)<br>8/8 (4-OHT) | N/A | 0/8 (TMX)<br>0/10 (4-OHT) |
| p5 | N/A | 7/8 (TMX) | N/A | 0/7 (TMX) |
| p16 | N/A | 9/9 (TMX) | 13/13 | 5/11 (TMX) |
| p21 | N/A | 8/8 (TMX) | N/A | 10/10 (TMX) |
| p26 | N/A | N/A | 6/6 | 6/6 (TMX) |
| P31 | N/A | 5/5 (TMX) | 6/6 | 7/7 (TMX) |
| p240 | N/A | 6/6 (TMX) | N/A | 7/7 (TMX) |

Figure S3

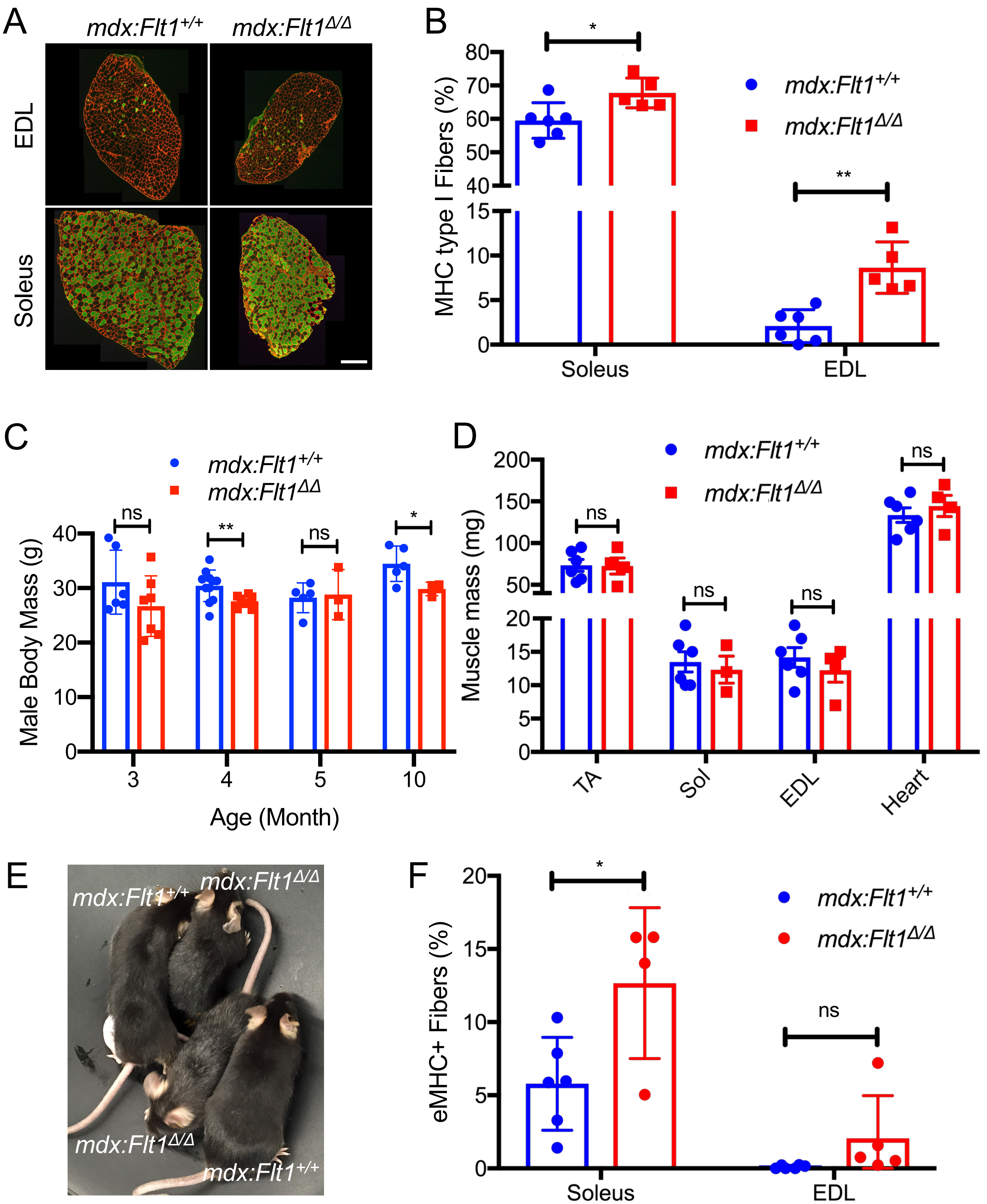
A
mdx:Flt1+/+
mdx:Flt1Δ/Δ
EDL
Soleus
B
MHC type I Fibers (%)
Soleus
EDL
C
Male Body Mass (g)
Age (Month)
D
Muscle mass (mg)
TA
Sol
EDL
Heart
E
F
eMHC+ Fibers (%)
ns
*
**

### Figure S4

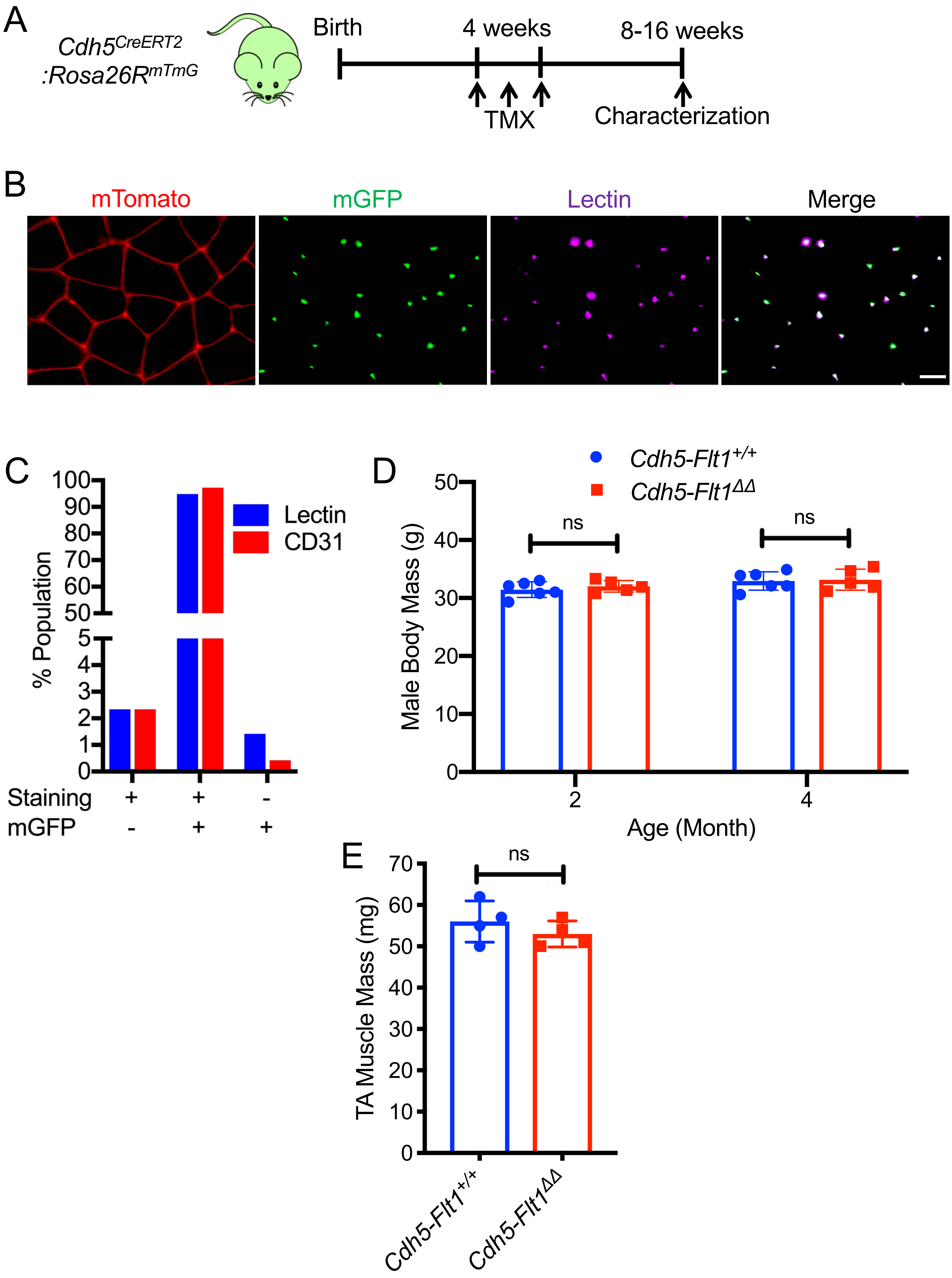
A
Cdh5^CreERT2 :Rosa26R^mTmG
Birth
4 weeks
8-16 weeks
TMX
Characterization
B
mTomato
mGFP
Lectin
Merge
C
% Population
Lectin
CD31
Staining + + -
mGFP - + +
D
Cdh5-Flt1+/+
Cdh5-Flt1ΔΔ
Male Body Mass (g)
ns
ns
Age (Month)
E
TA Muscle Mass (mg)
ns
Cdh5-Flt1+/+
Cdh5-Flt1ΔΔ

### Figure S5

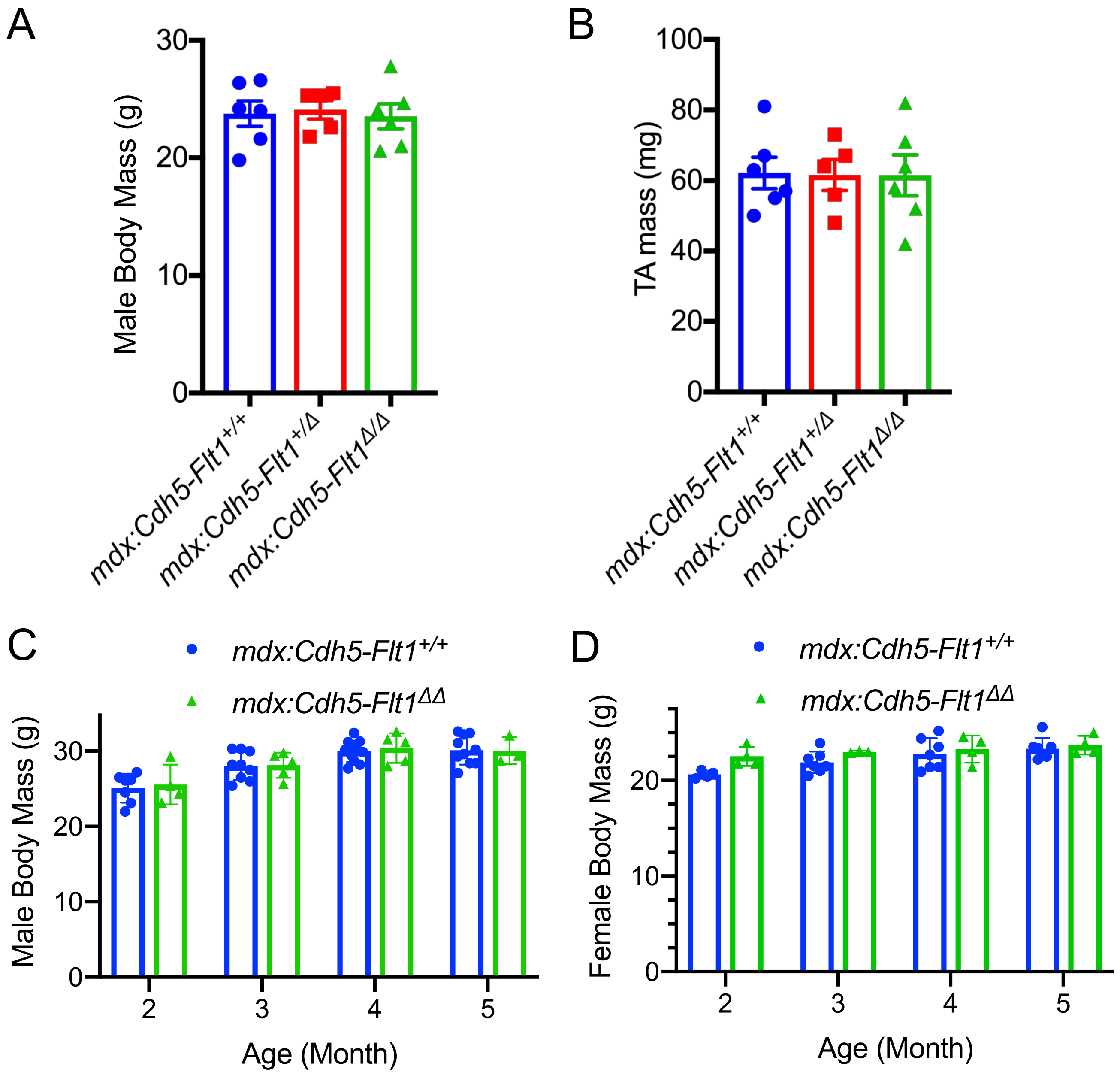

A
Male Body Mass (g)
30
20
10
0
mdx:Cdh5-Flt1+/+
mdx:Cdh5-Flt1+/Δ
mdx:Cdh5-Flt1Δ/Δ
B
TA mass (mg)
100
80
60
40
20
0
mdx:Cdh5-Flt1+/+
mdx:Cdh5-Flt1+/Δ
mdx:Cdh5-Flt1Δ/Δ
C
mdx:Cdh5-Flt1+/+
mdx:Cdh5-Flt1ΔΔ
Male Body Mass (g)
30
20
10
0
2
3
4
5
Age (Month)
D
mdx:Cdh5-Flt1+/+
mdx:Cdh5-Flt1ΔΔ
Female Body Mass (g)
20
10
0
2
3
4
5
Age (Month)

### Figure S6

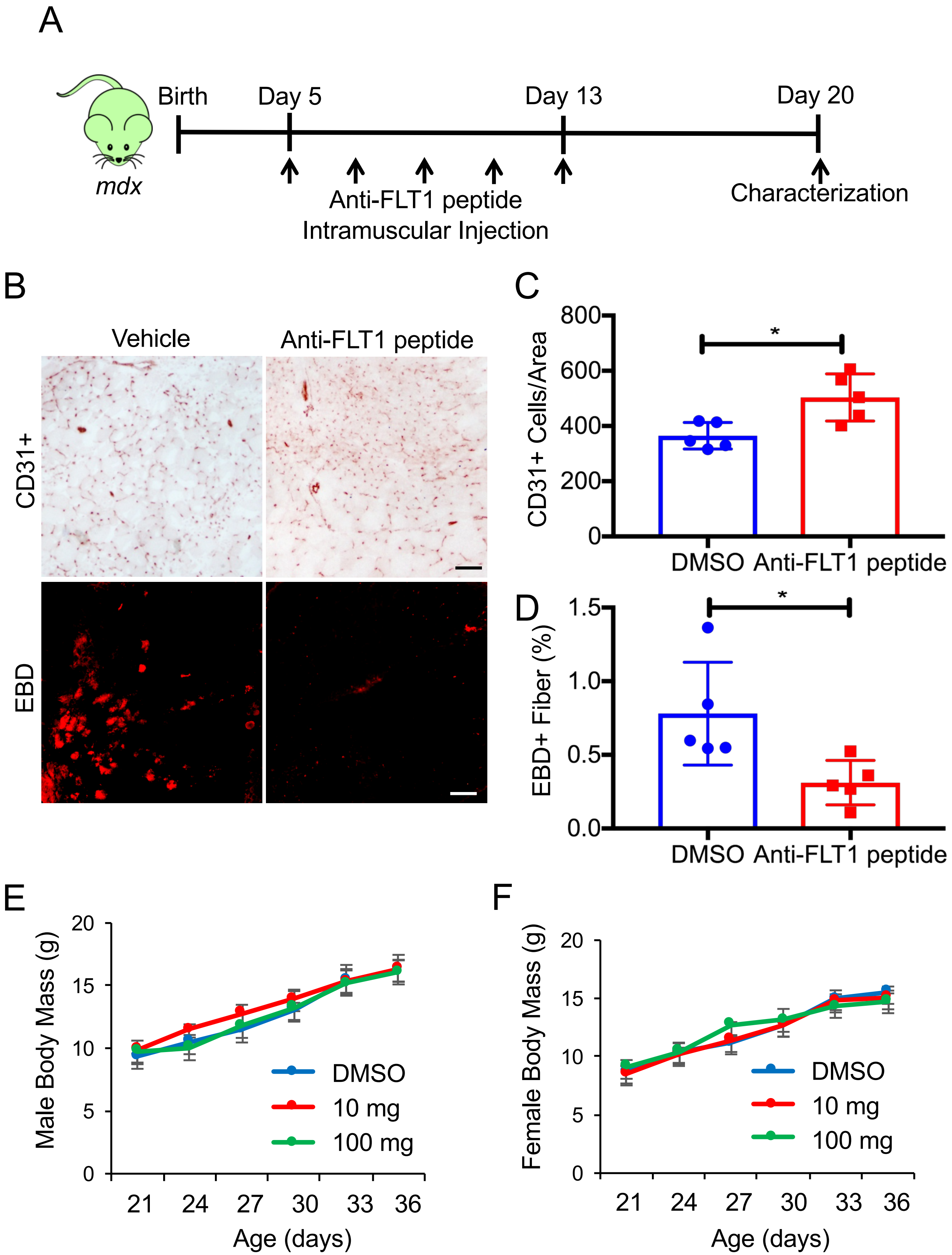
A
Birth
Day 5
Day 13
Day 20
mdx
Anti-FLT1 peptide
Intramuscular Injection
Characterization
B
Vehicle
Anti-FLT1 peptide
CD31+
EBD
C
CD31+ Cells/Area
800
600
400
200
0
*
DMSO
Anti-FLT1 peptide
D
EBD+ Fiber (%)
1.5
1.0
0.5
0.0
*
DMSO
Anti-FLT1 peptide
E
Male Body Mass (g)
20
15
10
5
0
21
24
27
30
33
36
Age (days)
DMSO
10 mg
100 mg
F
Female Body Mass (g)
20
15
10
5
0
21
24
27
30
33
36
Age (days)
DMSO
10 mg
100 mg

### Figure S7

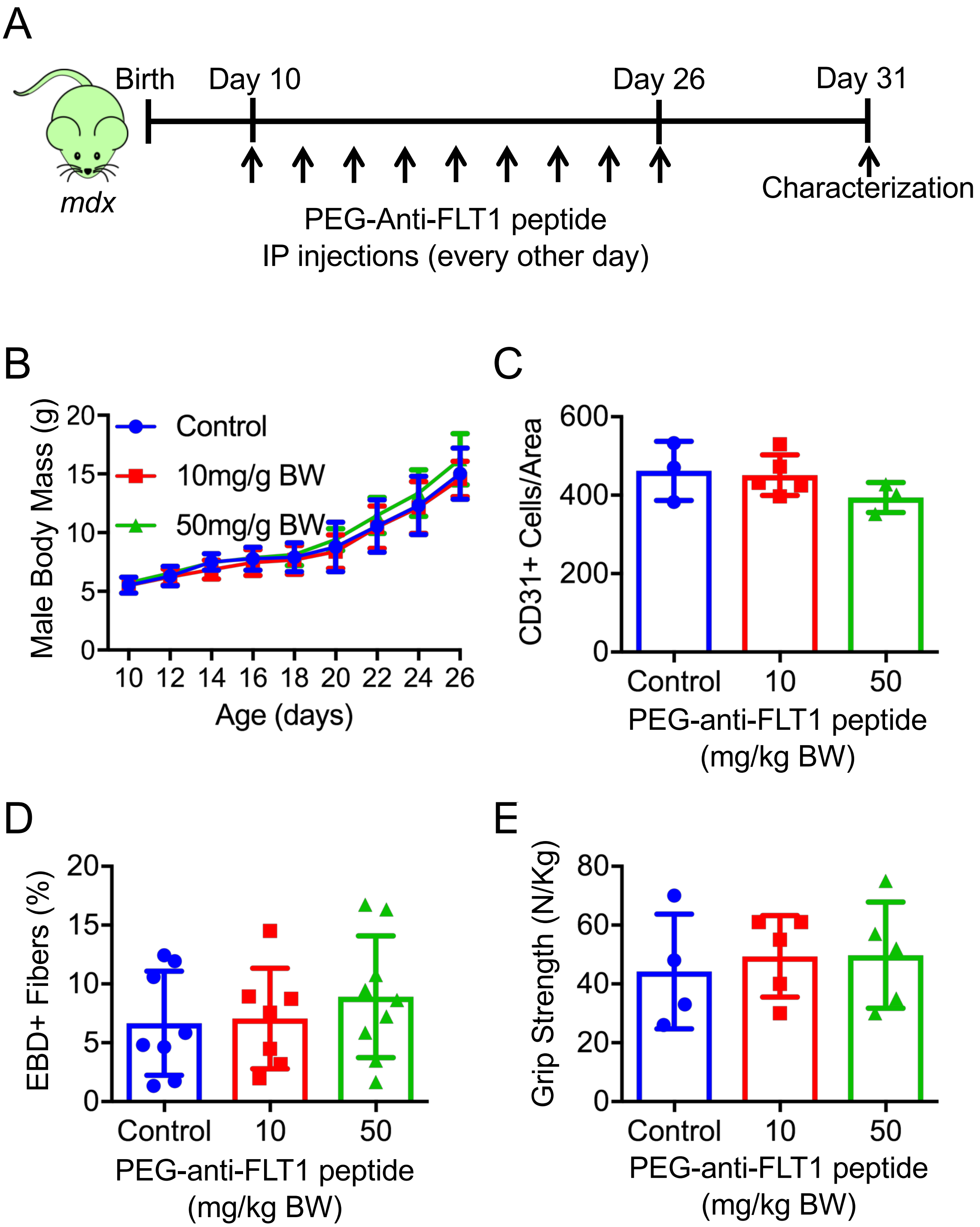

A
Birth
Day 10
Day 26
Day 31
mdx
Characterization
PEG-Anti-FLT1 peptide
IP injections (every other day)
B
Male Body Mass (g)
Control
10mg/g BW
50mg/g BW
Age (days)
C
CD31+ Cells/Area
Control
10
50
PEG-anti-FLT1 peptide
(mg/kg BW)
D
EBD+ Fibers (%)
Control
10
50
PEG-anti-FLT1 peptide
(mg/kg BW)
E
Grip Strength (N/Kg)
Control
10
50
PEG-anti-FLT1 peptide
(mg/kg BW)

### Figure S8

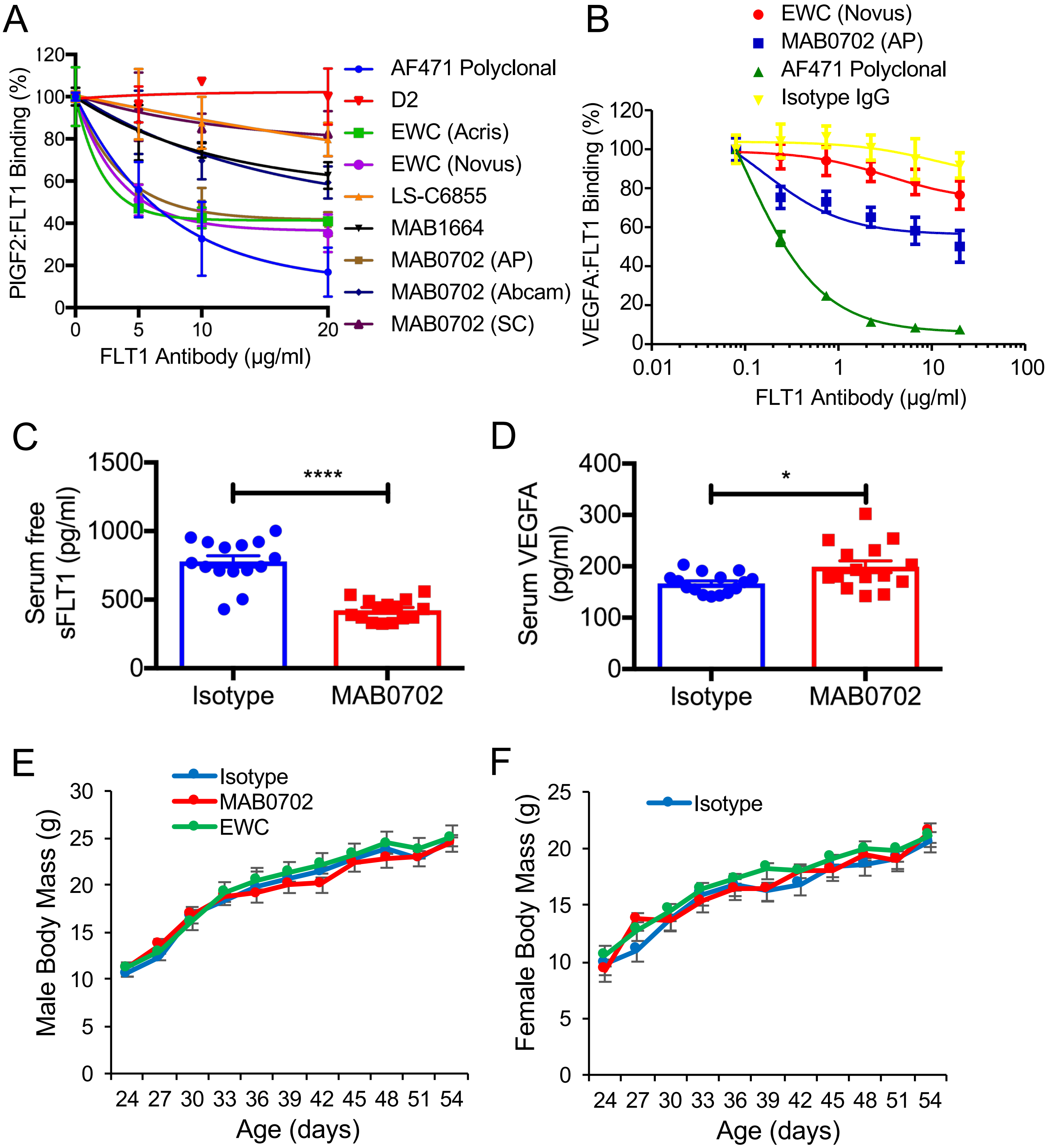
A
PlGF2:FLT1 Binding (%)
FLT1 Antibody (µg/ml)
AF471 Polyclonal
D2
EWC (Acris)
EWC (Novus)
LS-C6855
MAB1664
MAB0702 (AP)
MAB0702 (Abcam)
MAB0702 (SC)
B
EWC (Novus)
MAB0702 (AP)
AF471 Polyclonal
Isotype IgG
VEGFA:FLT1 Binding (%)
FLT1 Antibody (µg/ml)
C
Serum free sFLT1 (pg/ml)
****
Isotype
MAB0702
D
Serum VEGFA (pg/ml)
*
Isotype
MAB0702
E
Isotype
MAB0702
EWC
Male Body Mass (g)
Age (days)
F
Isotype
Female Body Mass (g)
Age (days)

### Table S1

| Table S2. Biacore analysis for affinity of monoclonal antibodies against FLT1 | | | |
|---|---|---|---|
| **Mouse FLT1** | **Ka ($M^{-1}s^{-1}$)** | **Kd ($s^{-1}$)** | **KD (M)** |
| MAB0702 | 5.555E+05 | 1.696E-03 | 3.053E-09 |
| EWC | 4.674E+05 | 3.488E-06 | 7.463E-12 |
| **Human FLT1** | **Ka ($M^{-1}s^{-1}$)** | **Kd ($s^{-1}$)** | **KD (M)** |
| MAB0702 | 3.811E+05 | 3.234E-04 | 8.486E-10 |
| EWC | 1.036E+05 | 6.625E-04 | 6.397E-09 |

### Table S2

**Table S1 DNA primer sequences**

| Gene | Forward | Sequence | Reverse | Sequence | Product (bp) |
|---|---|---|---|---|---|
| **Genotyping** | | | | | |
| *Flt1*$^{LoxP/LoxP}$ | Common | GTGCCACTGACCTAACATGTAAGAG | Common | GCGAAAAACAGTCAGTAGAAGT | 245 (WT)<br>280 (KI) |
| *CAG*$^{CreERTM}$ | OIMR1084 | GCGGTCTGGCAGTAAAAACTATC | OIMR1085 | GTGAAACAGCATTGCTGTCACTT | 102 (TG) |
| *Cdh5*$^{CreERT2}$ | CRE-1 | AACCTGGATAGTGAAACAGGGGC | ER-1 | CTCCATGGAGCGCCAGACGAGACC | 408 (TG) |
| **qPCR** | | | | | |
| *Flt1* | Exon 1 | CTTGCTCACCATGGTCAGCTGCTG | Exon 2 | CACTTTTAACTTCGACCCTGAGCC | 103 (WT/KO) |
| *Flt1* | Exon 2 | GGCCAGACTCTCTTTCTCAAGTGC | Exon 3 | GCAGAATTGCCTGTTATCCCTCCC | 135 (WT) |
| *18S rRNA* | 18S-F1 | CGCACGGCCGGTACAGTGAAACTG | 18S-R1 | CACCCGTGGTCACCATGGTAGGCA | 343 |
